# Epigenetic changes, neuronal dysregulation and metabolomic abnormalities in *Zmym2* mutant mice, a genetic model of schizophrenia and neurodevelopmental disorders

**DOI:** 10.1101/2025.02.18.638656

**Authors:** Wei-Chao Huang, Kira Perzel Mandell, Sameer Aryal, Bryan J. Song, Antia Valle-Tojeiro, Nathaniel Goble, Chuhan Geng, Ahmet S. Asan, Xiao-Man Liu, Courtney Dennis, Lucas Dailey, Amy Deik, Lucia Inunciaga, Eivgeni Mashin, Zohreh Farsi, Yining Wang, Jen Q. Pan, Clary B. Clish, Hasmik Keshishian, Steven A. Carr, Morgan Sheng

## Abstract

Loss-of-function mutations in *ZMYM2* are associated with an increased risk of schizophrenia (SCZ) and neurodevelopmental disorders (NDD). ZMYM2 interacts with proteins involved in histone modification and gene regulation, including LSD1 and ADNP; however, its specific roles in the brain remain poorly understood. In this multi-omics study, we demonstrate that heterozygous knockout of *Zmym2* in mice results in widespread disturbances in gene expression affecting diverse molecular pathways, including those related to histone modifications and neuronal activity. Proteomic analysis of synapses reveals dysregulation of lipid metabolism and neurofilament-associated pathways, while metabolomic profiling identifies alterations in sphingomyelin and ceramide levels. Furthermore, *Zmym2* mutant mice exhibit abnormal brain oscillation patterns on EEG and locomotor hyperactivity in the open field test. Collectively, these findings underscore the critical role of *ZMYM2* in brain development and function and highlight *Zmym2* mutant mice as a genetic animal model for SCZ and NDD.

## INTRODUCTION

Schizophrenia (SCZ) is a debilitating mental disorder with a poorly understood pathophysiology and limited treatment options, highlighting the need for deeper exploration of its underlying mechanisms. SCZ is strongly influenced by genetic factors, with heritability estimated at 60%- 80% (Sullivan et al., 2003). While genome-wide association studies (GWAS) have identified many common variants with small effects (Schizophrenia Working Group of the Psychiatric Genomics, 2014; Trubetskoy et al., 2022), recent exome sequencing studies, such as those from the Schizophrenia Exome Sequencing Meta-Analysis Consortium (SCHEMA), have discovered rare heterozygous loss-of-function (LoF) protein-truncating variants (PTVs) with a large impact on disease risk (Singh et al., 2022). The SCHEMA study (Singh et al., 2022), which analyzed 24,248 cases and 97,322 controls, identified 10 such risk genes with exome-wide significance and a total of 32 genes with a false discovery rate (FDR) of less than 5% for SCZ (hereafter referred to as SCHEMA genes).

Interestingly, a number of SCHEMA genes encode proteins involved in chromatin modification, transcription, or epigenetic regulation (Farsi and Sheng, 2023), including *ZMYM2* (zinc finger MYM-type containing 2, also known as ZNF198), which functions as a transcriptional corepressor (Connaughton et al., 2020). Heterozygous LoF mutations in *ZMYM2* have been linked to congenital anomalies of the kidney and urinary tract (CAKUT), with affected individuals also exhibiting neurological symptoms such as developmental delay (DD), intellectual disability (ID), autism spectrum disorder (ASD), and stereotypy (Connaughton et al., 2020). Furthermore, other independent studies have implicated *ZMYM2* variants in neurodevelopmental disorders (NDDs), ASD, and DD/ID (Ruzzo et al., 2019; Stessman et al., 2017; Wang et al., 2023a; Wang et al., 2020; Yuan et al., 2023), suggesting a critical role for *ZMYM2* in brain development and function.

ZMYM2 is an atypical zinc-finger protein whose MYM-type zinc finger is reported not to bind DNA (Gocke and Yu, 2008). However, it contains a multi-SUMO binding motif, enabling it to bind sumoylated proteins (Aguilar-Martinez et al., 2015). By recruiting SUMOylated HDAC1, ZMYM2 plays a crucial role in maintaining the LSD1-CoREST-HDAC1 (LCH) complex (Gocke and Yu, 2008), which functions as a histone H3K4 demethylase. LSD1 binds to DNA regulatory elements that highly overlap with the binding sites of SETD1A (Mukai et al., 2019), another high-confidence SCHEMA gene (Singh et al., 2022). Various studies have identified proteins that interact with ZMYM2 (Connaughton et al., 2020; Owen et al., 2023; Yang et al., 2020). Beyond the LCH complex, ZMYM2 binds to subunits of other complexes involved in transcriptional regulation, including ADNP and CHD4 (part of the ChAHP complex) (Kaaij et al., 2019); GTF3C1 and GTF3C4 (part of the TFIIIC complex) (Ferrari et al., 2020); ATF7IP and TRIM28 (part of the SETDB1–TRIM28 complex) (Tsusaka et al., 2020); and MGA (part of the PRC1.6 complex) (Mochizuki et al., 2021). This suggests that the loss of ZMYM2 could result in substantial transcriptional dysregulation. Indeed, deletion of ZMYM2 has been shown to reactivate silenced copies of imprinted genes—genes expressed from only one allele due to epigenetic silencing (Huang et al., 2018)—in human and mouse embryonic stem cells (Bar et al., 2021; Butz et al., 2022). Recent research has demonstrated that embryos with homozygous knockout of *Zmym2* (*Zmym2*^-/-^) exhibit de-repression of germline genes and long interspersed nuclear elements (LINEs), along with dysregulation of histone and DNA methylation (Graham-Paquin et al., 2023). Notably, many of the proteins that bind to ZMYM2 are encoded by risk genes for NDD and neuropsychiatric disorders. For example, *ADNP* is recognized as a high-confidence ASD risk gene (accounting for ∼0.17% of ASD cases) (Helsmoortel et al., 2014; Satterstrom et al., 2020), and mutations in *CHD4* can lead to DD/ID and speech delay (Weiss et al., 2020). Investigating the transcriptional dysregulations caused by *Zmym2* LoF may provide insights into the potential mechanisms underlying a variety of central nervous system (CNS) disorders.

To gain insights into the function of *ZMYM2* in the brain, we conducted transcriptomic profiling of multiple brain regions in *Zmym2* mutant mice at various ages, focusing on the heterozygous genotype that is particularly relevant to human disease. Additionally, we performed proteomic analysis of purified cortical synapses and metabolomic analysis of the cortex. Besides confirming the de-repression of germline genes, these comprehensive multi-omics investigations revealed unexpected molecular pathway changes in *Zmym2* heterozygous (*Zmym2*^+/-^) brains, such as alterations in activity-regulated genes (ARGs) in neurons. Furthermore, *Zmym2*^+/-^ mouse mutants showed abnormal brain oscillations by EEG and hyperactive locomotor behavior. Our study provides a comprehensive characterization of molecular, cellular, and systems-level changes in *Zmym2* mutant mice, contributing to a deeper mechanistic understanding of SCZ and NDD.

## RESULTS

### Widespread dysregulation of gene expression, including de-repression of germline genes and LINE1 retrotransposon elements in the *Zmym2*^+/-^ mutant brain

In breeding of *Zmym2*^+/-^ mice, no *Zmym2*^-/-^ pups were born (Supplemental Fig. 1A), consistent with previous reports of embryonic lethality in *Zmym2*^-/-^ mice (Graham-Paquin et al., 2023). Consequently, our study focuses on *Zmym2*^+/-^ mice, which is also the most relevant genotype for SCZ risk.

*Zmym2*^+/-^ brain exhibited ∼50% reduction in *Zmym2* RNA and ZMYM2 protein levels, compared with wildtype (WT) littermate controls (Supplemental Fig. 1B, C). While body lengths were unchanged, the body and brain weights of *Zmym2*^+/-^ mice were significantly lower than those of WT mice (by ∼5-10%) (Supplemental Fig. 1D-F), which is interesting given that *ZMYM2* and its binding partner *ZMYM3* have been implicated in DD and microcephaly (Connaughton et al., 2020; Hiatt et al., 2023).

To investigate gene expression changes across different brain regions and ages, we performed bulk messenger RNA sequencing (RNA-seq) of prefrontal cortex (PFC), hippocampus (HP), somatosensory cortex (SSC), striatum (STR), thalamus (TH), and substantia nigra (SN) on *Zmym2*^+/-^ and WT male mice at 1, 3, and 5 months old of age (1-, 3-, and 5-mo). A large number of differentially expressed genes (DEGs, defined by FDR-adjusted p value [padj] < 0.05) were observed in all examined brain regions and ages. In general, there were more upregulated DEGs than downregulated DEGs across brain regions and developmental stages, except for 3-mo STR and SN (Fig. 1A, Supplemental table 1). This pattern is consistent with ZMYM2’s role as part of a transcriptional corepressor complex (Connaughton et al., 2020).

**Figure 1:**
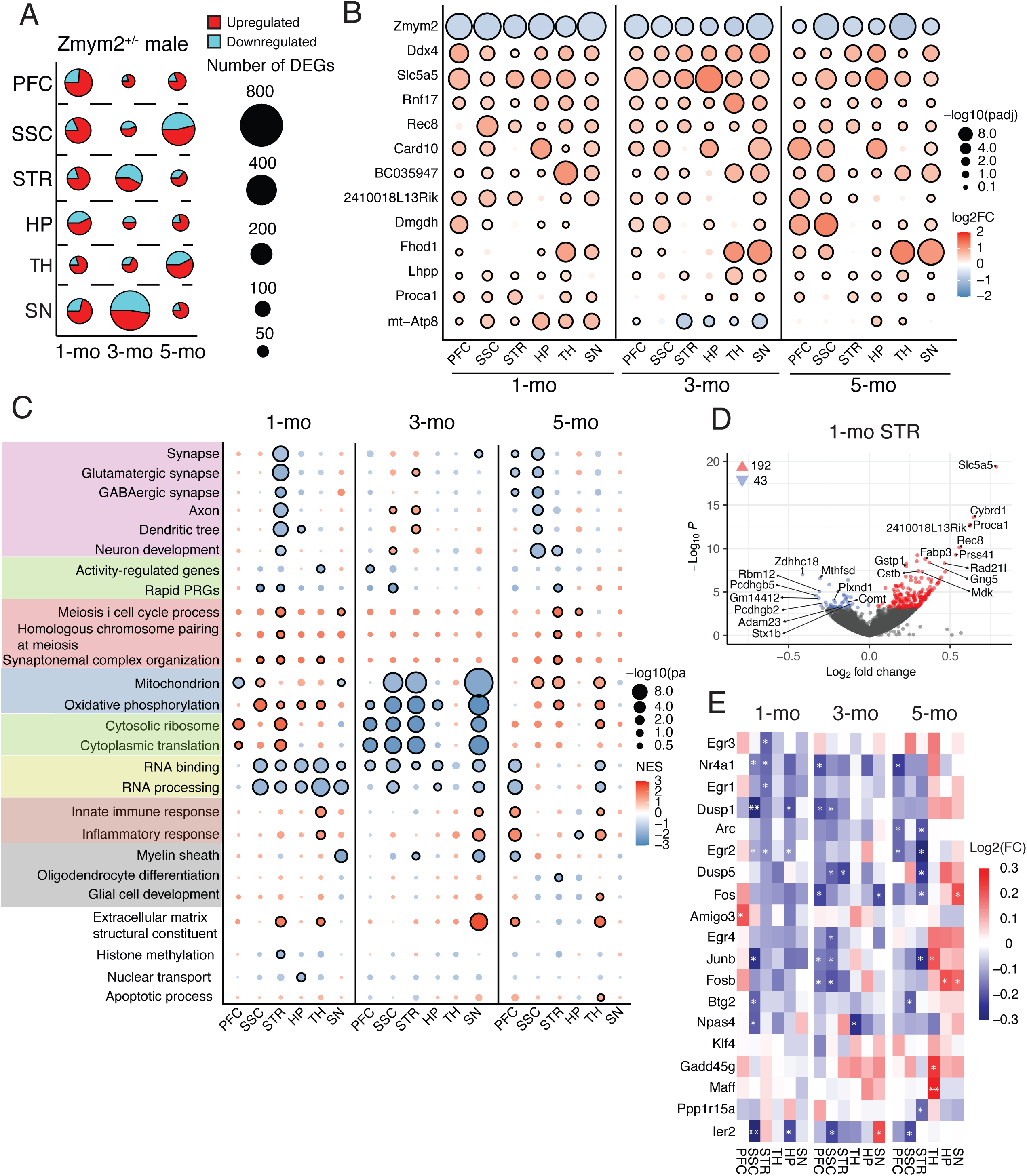
Transcriptomic changes in multiple molecular pathways in the *Zmym2*^+/−^ brain (A) Number of DEGs in each indicated brain region and age in *Zmym2* mutant mice. The size of the circles represents the number of DEGs, defined as those with a padj < 0.05, while the color of the circles indicates the direction of expression changes. (B) Bulk RNA-seq expression changes in *Zmym2*^+/-^ mice across brain regions and ages. Statistical significance is represented by circle size, and log2FC by the color scale. Circles with black outlines indicate padj < 0.05. (C) GSEA results of transcriptomic changes in *Zmym2*^+/-^ mice at 1-, 3-, and 5-mo across brain regions and developmental stages. Statistical significance is represented by circle size, and normalized enrichment score (NES) by the color scale. Circles with black outlines indicate an FDR < 0.05. (D) Volcano plots of bulk RNA-seq transcriptomic changes in the 1-mo STR of *Zmym2*^+/-^ mice. Red and blue dots represent DEGs (padj < 0.05). The numbers indicate the number of upregulated (red arrowhead) and downregulated (blue arrowhead) DEGs. (E) Heatmap showing log2FC values of rapid PRGs from bulk RNA-seq data in the indicated brain regions of *Zmym2*^+/-^ mice. *: p < 0.05; **: padj < 0.05.

To understand whether the observed DEGs overlap across brain regions, we compared the DEGs at each age and found only a small number that were common across the six examined brain regions at each age (1-mo: 12; 3-mo: 5; 5-mo: 7 DEGs, Supplemental Fig. 2A). We found 12 DEGs with significant differential expression changes in more than 12 RNA-seq brain regions out of a total of 18 (across 3 ages) (Fig. 1B). Interestingly, some of these 12 genes had been also identified as upregulated DEGs in *Zmym2*^-/-^ embryos (Graham-Paquin et al., 2023), for example, *Ddx4* and *Rnf17* (Pan et al., 2005; Tanaka et al., 2000), which are highly expressed in the testis and play important roles in spermatogenesis, suggesting that *Zmym2* heterozygous LoF causes the de-repression of germline genes in the brain.

More broadly than just looking at common DEGs, we asked whether *Zmym2* mutation leads to similar gene expression alterations across different brain regions. Indeed, a transcriptome-wide comparison revealed modest, positive correlations in log2-fold change (log2FC) values of individual genes across the examined brain regions at the same developmental stage (Spearman’s r: 0.18–0.35 at 1-mo; 0.16-0.33 at 3-mo; 0.2-0.48 at 5-mo; Supplemental Fig. 2B). These correlation coefficients between different brain regions of the same age were notably higher than those observed in the *Grin2a* mouse model of SCZ (Farsi et al., 2023) (Supplemental Fig. 2C).

To gain insight into the biological pathways altered in *Zmym2* mutant brain, we performed gene set enrichment analysis (GSEA) of bulk transcriptomic data. Numerous gene ontology (GO) terms were found to be significantly (FDR < 0.05) enriched among upregulated and downregulated genes (hereafter referred to as upregulated or downregulated GO terms/gene sets) across multiple brain regions. Notably, meiosis-related GO terms were consistently upregulated in *Zmym2* mutant mice, with the strongest enrichment observed in the STR (Fig. 1C, Supplemental Table 2). In addition, we detected upregulation of L1MdA, an active murine-specific LINE-1 transposable element (Sookdeo et al., 2013), in many brain regions at 1-mo, followed by downregulation at 3-mo (Supplemental Fig. 2D). Previous studies reported de-repression of germline genes and LINE elements in *Zmym2*^-/-^ embryos (Graham-Paquin et al., 2023). While our findings on germline gene regulation align with observations in early development, the transient upregulation of LINE-1 elements in the postnatal brain suggests that ZMYM2 plays a dynamic role in gene regulation.

### Differential alterations of diverse molecular pathways across brain regions of *Zmym2* mutants

Besides germline genes, GSEA identified several other biological processes altered in *Zmym2* mutant brains (Fig. 1C). These included GO terms related to synapses, ribosome/translation, and mitochondria/oxidative phosphorylation (OXPHOS).

Synapse-related gene sets, including those for glutamatergic and GABAergic synapses, were significantly downregulated in several brain regions, with an interesting age-dependence (Fig. 1C). In juvenile (1-mo) mice, synapse-related terms were downregulated primarily in the STR (Fig. 1C), but this effect diminished by 3-and 5-mo. Instead, synapse downregulation shifted to the PFC and SSC at 5-mo (Fig. 1C).

Consistent with GSEA, multiple synapse-related genes were significantly downregulated in 1-mo *Zmym2*^+/-^ mice in the STR, a brain region implicated in SCZ pathophysiology (McCutcheon et al., 2019). These included *Plxnd1* (a semaphorin receptor regulating striatal synapse formation) (Ding et al., 2011), *Stx1b* (Syntaxin 1B; an epilepsy risk gene) (Wolking et al., 2019), and *Comt* (an enzyme involved in dopamine metabolism, which has also been shown to be suppressed in *Grin2a* mutant mice) (Farsi et al., 2023) (Fig. 1D).

To probe more deeply into the synapse-related transcriptomic changes in *Zmym2* mutants, we performed GSEA with refined SynGO terms (Koopmans et al., 2019). By SynGO analysis, synapse-related gene sets were again most prominently changed (downregulated) in 1-mo STR of *Zmym2*^+/-^ mutant mice, affecting both pre-and postsynaptic processes (Supplemental Fig. 3A). At 1-mo, upregulation of SynGO terms for “translation at presynapse” and “translation at postsynapse” (Supplemental Fig. 3A) was observed especially in PFC, SSC, and STR, mirroring the ribosome/translation-related changes observed in GSEA with standard GO terms (Fig. 1C). Notably, the ribosome/translation-related pathways, including translation at presynapse and postsynapse, reversed direction (became downregulated) at 3-mo (Fig. 1C, Supplemental Fig. 3A); and in keeping with this, the expression of genes encoding cytosolic ribosome subunits showed a negative correlation between 1-and 3-mo in *Zmym2*^+/-^ mice (Supplemental Fig. 3B). Mitochondria/OXPHOS terms changed in generally the same direction as ribosome/translation terms in *Zmym2* mutant brain (Fig. 1C, Supplemental Fig. 3B, C), perhaps reflecting coordinated changes in cellular energetics and metabolism in different brain regions at various stages of brain maturation.

How is neuronal activity in different brain regions affected by *Zmym2* LoF? We used expression of known ARGs as a surrogate measure of neural activity. GSEA of bulk RNA-seq data showed that rapid primary response genes (PRGs), a set of genes that are induced quickly upon neuronal activation (Tyssowski et al., 2018), were downregulated in several brain regions, especially in the SSC and STR at 1-mo (Fig. 1C). For example, expression of well-known immediate early genes (IEGs) such as *Fos*, *Arc*, *Nr4a1* and *Egr1* (Okuno, 2011), was reduced in most examined brain regions at 1-and 3-mo (Fig. 1E).

We next asked whether the transcriptomic changes of *Zmym2*^+/-^ mice were enriched for genes associated with SCZ and/or other neuropsychiatric disorders. Genes associated with NDD were significantly enriched among downregulated genes in multiple brain regions including STR (Supplemental Fig. 3D), consistent with the known association of *ZMYM2* with NDD (Wang et al., 2020). SCHEMA genes (rare variant SCZ risk genes) (Singh et al., 2022) were significantly enriched particularly among downregulated genes in the STR at 1-and 5-mo (Supplemental Fig. 3D).

We also performed RNA-seq on 1-mo female mice to look for potential sex differences in *Zmym2* mutant mice. Similar to males, the majority of DEGs in females were upregulated rather than downregulated (Supplemental Fig. 4A). Remarkably, of the 12 DEGs that were upregulated across most brain regions in the male *Zmym2*^+/-^ (Fig. 1B), the vast majority also exhibited upregulation in 1-mo *Zmym2*^+/-^ females (Supplemental Fig. 4B). GSEA revealed that, as observed in males, meiosis-related pathways were upregulated in *Zmym2*^+/-^ females (Supplemental Fig. 4C, D, Supplemental Table 2). However, *Zmym2* LoF resulted in distinct alterations in other molecular pathways in 1-mo females (Supplemental Fig. 4C). Specifically, *Zmym2*^+/-^ females showed a decrease in synapse-and myelin-related pathways in PFC, whereas male mutants did not (Supplemental Fig. 4C).

### Transcriptomic changes in neuronal and non-neuronal cell types in *Zmym2* mutant mice

To investigate the impact of *Zmym2* LoF on specific cell types, we performed single-nucleus RNA-seq (snRNA-seq) analysis in PFC (1-and 5-mo) and STR (1-mo). No significant differences were observed in the proportions of major cell types between *Zmym2*^+/-^ and WT mice (Supplemental Fig. 5A, Supplemental table 3).

Differential expression analysis of snRNA-seq data revealed a sizable number of DEGs (defined as FDR < 0.05) in excitatory neurons of the PFC at 1-and 5-mo, and in inhibitory spiny projection neurons (SPNs) of the 1-mo STR (Fig. 2A, Supplemental table 3). Both excitatory and inhibitory neurons exhibited predominantly upregulated DEGs (Fig. 2A). Many of the upregulated meiosis-related DEGs, such as *Rnf17* and *Rec8*, were consistent between bulk RNA-seq (Fig. 1B) and snRNA-seq (Fig. 2B). *Zmym2* is expressed in neurons and glial cells, with higher expression in neurons (http://dropviz.org/). All major glial cell types, including astrocytes, oligodendrocytes, and oligodendrocyte progenitor cells (OPCs), were affected by *Zmym2* LoF, particularly in the 1-mo STR, where oligodendrocytes and astrocytes had more DEGs than the inhibitory neurons (Fig. 2A).

**Figure 2:**
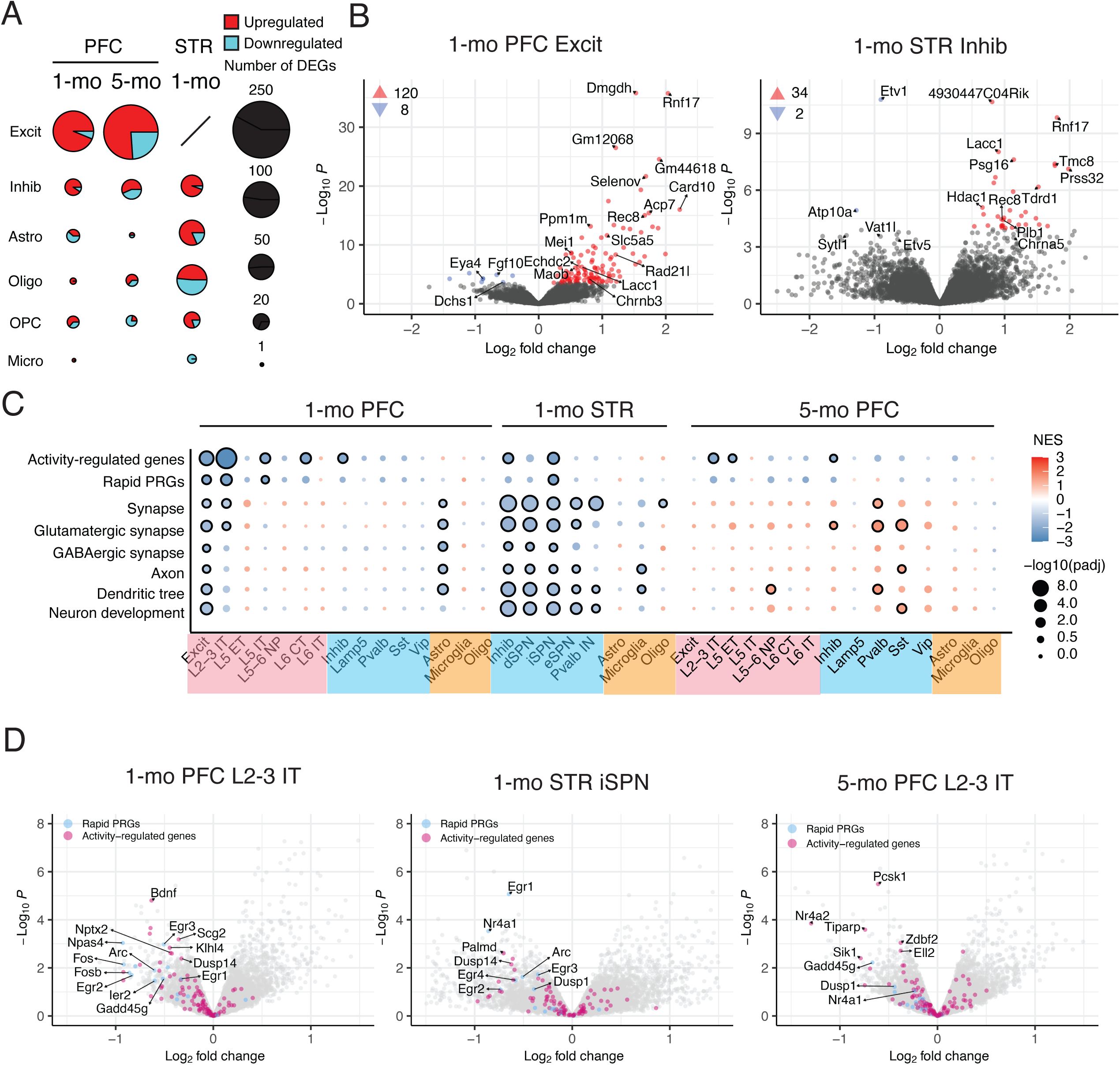
Dysregulation of activity-and synapse-related pathways in neuronal cells of *Zmym2*^+/-^ mice (A) Number of DEGs (FDR < 0.05) across cell types in the indicated brain regions, with circle color indicating the direction of expression change. Excit: excitatory neuron; Inhib: inhibitory neuron; Astro: astrocyte; Oligo: oligodendrocyte; OPC: oligodendrocyte progenitor cells; Micro: microglia. No DEGs were identified in microglia in the 5-mo PFC. Excitatory neurons were not identified in the 1-mo STR. (B) Volcano plots of transcriptomic changes from snRNA-seq analysis in specific cell types and brain regions. Red and blue dots represent genes with FDR < 0.05. The numbers represent the number of upregulated and downregulated DEGs. (C) GSEA results for indicated GO terms in brain regions of *Zmym2*^+/-^ mice. L2–L6: cortical layers 2–6; IT: intratelencephalic; CT: corticothalamic neurons; SST: somatostatin interneurons; VIP: vasoactive intestinal peptide interneurons; Lamp5: Lamp5 GABAergic cortical interneuron; dSPN: direct-pathway spiny projection neuron; iSPN: indirect-pathway SPN; eSPN: eccentric SPN; Pvalb: parvalbumin interneurons. (D-F) Volcano plots of transcriptomic changes in neuron subtypes in the indicated brain regions of *Zmym2*^+/-^ mice, highlighting ARGs (red dots) and rapid PRGs (blue dots).

GSEA revealed upregulation of meiosis-related GO terms predominantly in neuronal cell types in PFC and STR (Supplemental Fig. 5B, Supplemental table 4). Downregulation of ARGs and rapid PRGs, in PFC and STR, occurred mainly in neuronal cell types – both excitatory and inhibitory (Fig. 2C). Reduced expression of ARGs, including possible SCZ biomarkers *Bdnf* (Mohammadi et al., 2018) and *Nptx2* (Aryal et al., 2023; Kimoto et al., 2015; Xiao et al., 2021), as well as prominent downregulation of rapid PRGs such as *Egr1*, *Arc*, and *Nr4a1*, was observed across various subtypes of excitatory and inhibitory neurons, particularly in the PFC (e.g., L2-3 IT neurons) and STR (SPNs) (Fig. 2C, D). In addition, GO terms related to synapse were significantly downregulated in excitatory neurons of PFC and inhibitory neurons of STR at 1-mo (Fig. 2C). Interestingly, synapse and glutamatergic synapse-related terms were upregulated at 5-mo in PFC, most prominently in parvalbumin and SST inhibitory interneurons, suggesting some synaptic dysregulation or compensation in these interneurons, which are implicated in SCZ pathophysiology (Dienel and Lewis, 2019; Duncan et al., 2025).

GO terms related to steroid/cholesterol biosynthesis were upregulated in astrocytes of 1-mo PFC of *Zmym2*^+/-^ mice (Supplemental Fig. 5B, C), which is reminiscent of similar changes in astrocytes of *Grin2a* mutant mice (Farsi et al., 2023), but in the opposite direction to changes in astrocytes noted in human SCZ postmortem brain tissues (Ling et al., 2024). The upregulation of steroid synthesis pathways in astrocytes of PFC was associated with upregulation of ribosome/translation and mitochondria/OXPHOS gene sets in astrocytes (Supplemental Fig. 5B) and with downregulation of synapse-related and ARG gene sets in neurons of PFC (Fig. 2C).

### *Zmym2* LoF dysregulates epigenetic pathways in the brain

*Zmym2* LoF de-represses germline genes and retrotransposon elements, a process linked to H3K4 hypermethylation and loss of DNA methylation (Graham-Paquin et al., 2023). Furthermore, ZMYM2 interacts with several subunits of epigenetic regulatory complexes. Given these roles, we investigated whether *Zmym2* LoF leads to alterations in epigenetic pathways in the brain.

GSEA of bulk RNA-seq data revealed significant downregulation of pathways related to histone modification and chromatin remodeling, particularly in the STR (1-and 5-mo) and the TH (3-and 5-mo) (Fig. 3A, B). Many of the specific downregulated genes in these pathways are associated with NDDs (Fig. 3B). For instance, *Suv39h2*, a lysine-specific methyltransferase involved in histone modification, has been implicated in ASD (Balan et al., 2021); similarly, *Kmt2a*, another lysine-specific methyltransferase, is a causative gene in Wiedemann-Steiner syndrome (Jones et al., 2012). The “SWI/SNF superfamily type complex” and the “RNAi effector complex” were both downregulated in the STR of *Zmym2* mutant mice (Fig 3A, B). The SWI/SNF complex plays a critical role in chromatin remodeling, and variants in multiple SWI/SNF components have been reported to cause ID (Bogershausen and Wollnik, 2018). The Argonaute gene family (e.g., *Ago1* & *Ago2*), essential for RNAi, is implicated in NDDs, collectively referred to as Argonaute syndromes (https://argonautes.ngo/en). Notably, *Hdac1* (Histone Deacetylase 1; an enzyme that removes acetyl groups from lysine residues on histones) was increased in the bulk RNA-seq analysis of the STR in 1-mo male (log2FC = 0.23, padj = 0.056) and female (log2FC =0.26, padj = 0.026) *Zmym2*^+/-^ mice (Fig. 3C), and *Hdac1* mRNA was increased in most examined brain regions of *Zmym2* mutants (Fig. 3C). In snRNA-seq, *Hdac1* was also significantly upregulated in inhibitory neurons of STR (Fig. 2B). By western blotting, we found ∼50% increase in HDAC1 protein levels in 1-mo STR of *Zmym2*^+/-^ mice (Fig. 3D).

**Figure 3:**
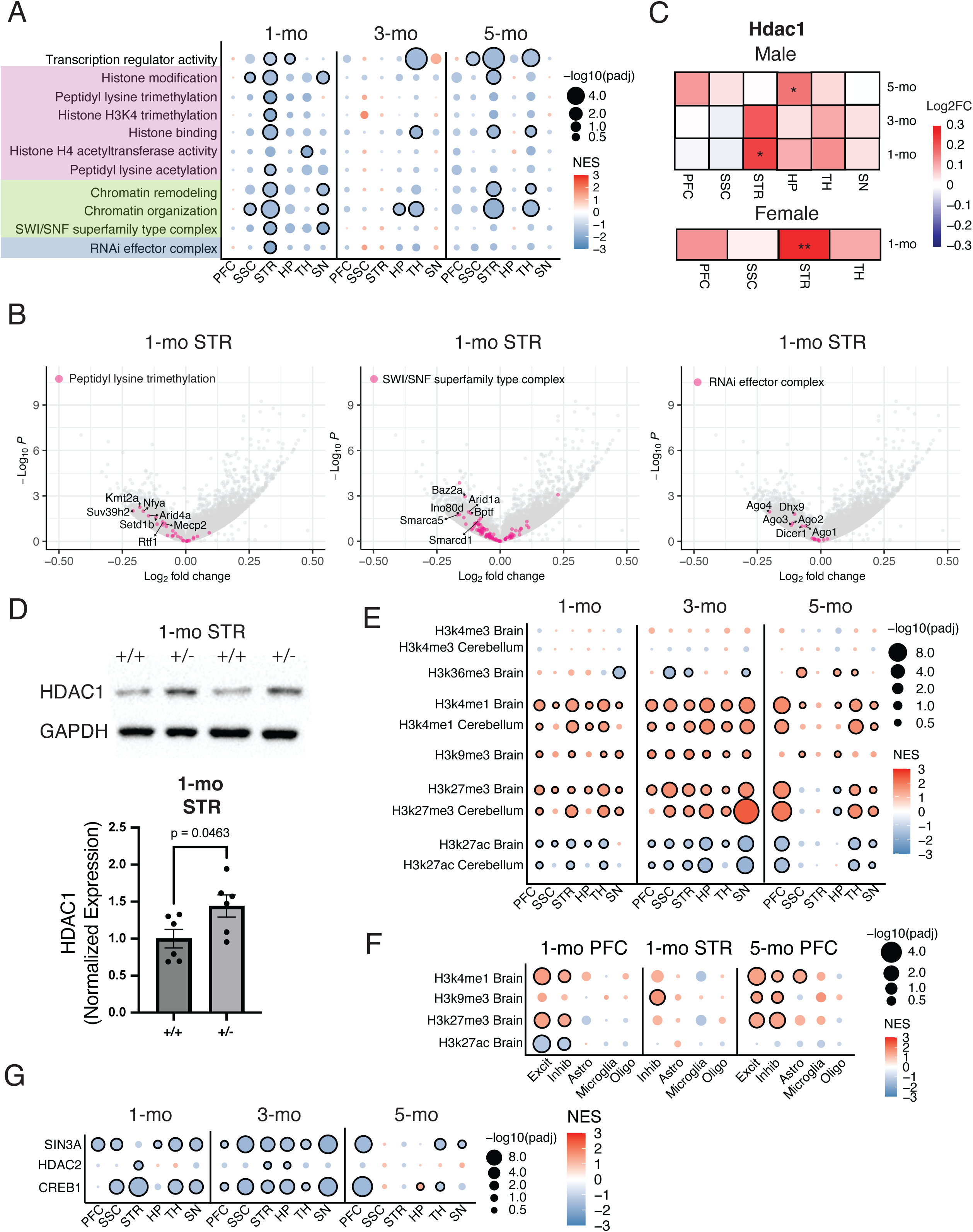
Dysregulation of epigenetic pathways in *Zmym2*^+/–^ mutant mice (A) GSEA results in indicated brain regions of *Zmym2*^+/-^ mice. (B) Volcano plots of transcriptomic changes in 1-mo *Zmym2*^+/-^ STR, highlighting genes from the indicated gene sets. (C) Heatmap showing log2FC values of *Hdac1* in bulk RNA-seq data from the indicated brain regions in male and female *Zmym2* mutant mice. *: p < 0.05; **: padj < 0.05 (D) Western blot analysis of HDAC1 and GAPDH in total striatal lysates from 1-mo *Zmym2* males. Bar plots show the quantification of HDAC1 protein levels in the STR of *Zmym2* animals (n = 5-6 per genotype). Error bars represent the mean ± standard error. Each chart indicates the p value from a two-tailed t-test. (E-F) GSEA results for genes marked by histone modifications from bulk and snRNA-seq. (G) GSEA results for genes bound by transcription factors from bulk RNA-seq.

We further analyzed our bulk RNA-seq results against chromatin immunoprecipitation (ChIP) sequencing data from the ENCODE database (Consortium, 2012) to assess whether genes marked by histone modifications were enriched among differentially regulated genes in the *Zmym2*^+/-^ brain. Consistent with previous findings that *Zmym2* knockout causes upregulation of H3K4me1 levels (Yang et al., 2020), an active marker of gene expression, upregulated genes in multiple brain regions of *Zmym2*^+/-^ were significantly enriched for H3K4me1-marked genes (Fig. 3E). Genes marked with H3K9me3, a repressive marker methylated by the complex to which ZMYM2 binds (Tsusaka et al., 2020), were also enriched among upregulated genes in *Zmym2*^+/-^ mice, suggesting that these genes are de-repressed in these mutants. Additionally, H3K27me3-marked genes were enriched among upregulated genes, whereas H3K27ac-marked genes were enriched among downregulated genes of *Zmym2*^+/-^ mice. The snRNA-seq analysis revealed enrichment patterns similar to those observed in the bulk RNA-seq data (Fig. 3F). These GSEA analyses provide evidence that *Zmym2* LoF affects gene expression regulated through histone modifications and offer insights into which histone modifications are potentially influenced by *Zmym2* LoF.

In addition to histone modifications, we examined the enrichment of transcription factor-bound genes among differentially regulated genes in *Zmym2*^+/-^ brains using publicly available ChIP-seq datasets (Consortium, 2012; Keenan et al., 2019). SIN3A forms a histone deacetylase complex with HDAC1/2 and other associated proteins, functioning as a transcriptional repressor (Kadamb et al., 2013). Loss of SIN3A has been shown to enhance synaptic potentiation and increase the expression of synaptic genes (Bridi et al., 2020), suggesting that the SIN3A-HDAC complex plays a role in repressing synaptic gene expression and neuronal plasticity. In *Zmym2*^+/-^ mice, SIN3A-bound genes were significantly enriched among downregulated genes, along with those bound by HDAC2, across multiple brain regions (Fig. 3G). Similarly, genes bound by CREB1, a transcription factor that recruits the histone acetyltransferase CBP/p300 (CREB-binding protein) and regulates numerous ARGs (Sakamoto et al., 2011; Tullai et al., 2007), were enriched among downregulated genes (Fig. 3G). These findings suggest that histone deacetylation mediated by the HDAC1-containing complex may contribute to transcriptional downregulation in *Zmym2*^+/-^ brains.

### *Adnp* and *Zmym2* LoF show similar transcriptomic changes

A previous study demonstrated that *ADNP* and *POGZ*, which encode proteins known to interact with ZMYM2 (Owen et al., 2023), are high-confidence ASD risk genes (Satterstrom et al., 2020). ADNP syndrome (OMIM #615873), White-Sutton syndrome (caused by heterozygous mutation in the *POGZ* gene; OMIM #616364), and ZMYM2 mutation-associated disorder (OMIM #619522) share overlapping clinical features, including hypotonia, DD/ID, and ASD traits. To determine whether LoF of these genes leads to similar transcriptome profiles, we compared the transcriptomic data from *Zmym2*^+/-^ mutants with publicly available RNA-seq results from the brains of *Adnp* (Cho et al., 2023) and *Pogz* (Suliman-Lavie et al., 2020) mutant mouse models.

We first identified the nominally significant (p value < 0.05) DEGs that were shared across the brain regions from *Adnp*, *Pogz*, and *Zmym2* mutants (*Adnp*: juvenile HP [AJ-HP] & adult HP [AA-HP]; *Pogz*: adult HP [P-HP] & adult cerebellum [P-Cere]; *Zmym2*: 1-mo & 3-mo HP). This analysis revealed 4 genes shared across all six regions and 42 genes shared in only adult HP (Supplemental Fig. 6A, B). When focusing on the set of consistently significant DEGs in *Zmym2*^+/-^ mice (Fig. 1B), we observed that they were mostly upregulated in *Adnp* mutants but downregulated in *Pogz* (Supplemental Fig. 6C).

Additionally, we performed GSEA on RNA-seq data from *Adnp* and *Pogz* mutant mice and compared them with *Zmym2^+/-^* mice. Among the altered GO terms (p < 0.05) shared across all three mouse mutants, *Adnp* and *Zmym2* exhibited the same directionality in most of the GO terms, contrasting with *Pogz* (Supplemental Fig. 6D, E). Thus, *Zmym2* mutant mice resemble *Adnp* but not *Pogz* mutant mice, in terms of transcriptomic changes in the brain.

### Changes in the synaptic proteome of *Zmym2* mutant mice

To understand the impact of *Zmym2* heterozygous LoF on synapses at the protein level, we conducted quantitative mass spectrometry (MS)-based proteomics on synaptic fractions purified from the cerebral cortex of 1-and 3-mo *Zmym2*^+/-^ mice and their WT littermates. Numerous differentially expressed proteins (DEPs, p < 0.01) were identified in synaptic proteomics of *Zmym2*^+/-^ mice at both ages (Fig. 4A, Supplemental table 5). The DEPs showed a moderate positive correlation between 1-& 3-mo (Spearman r = 0.48) (Supplemental Fig. 7A); the correlation of all protein changes between the two ages was weaker (Spearman r = 0.18).

**Figure 4:**
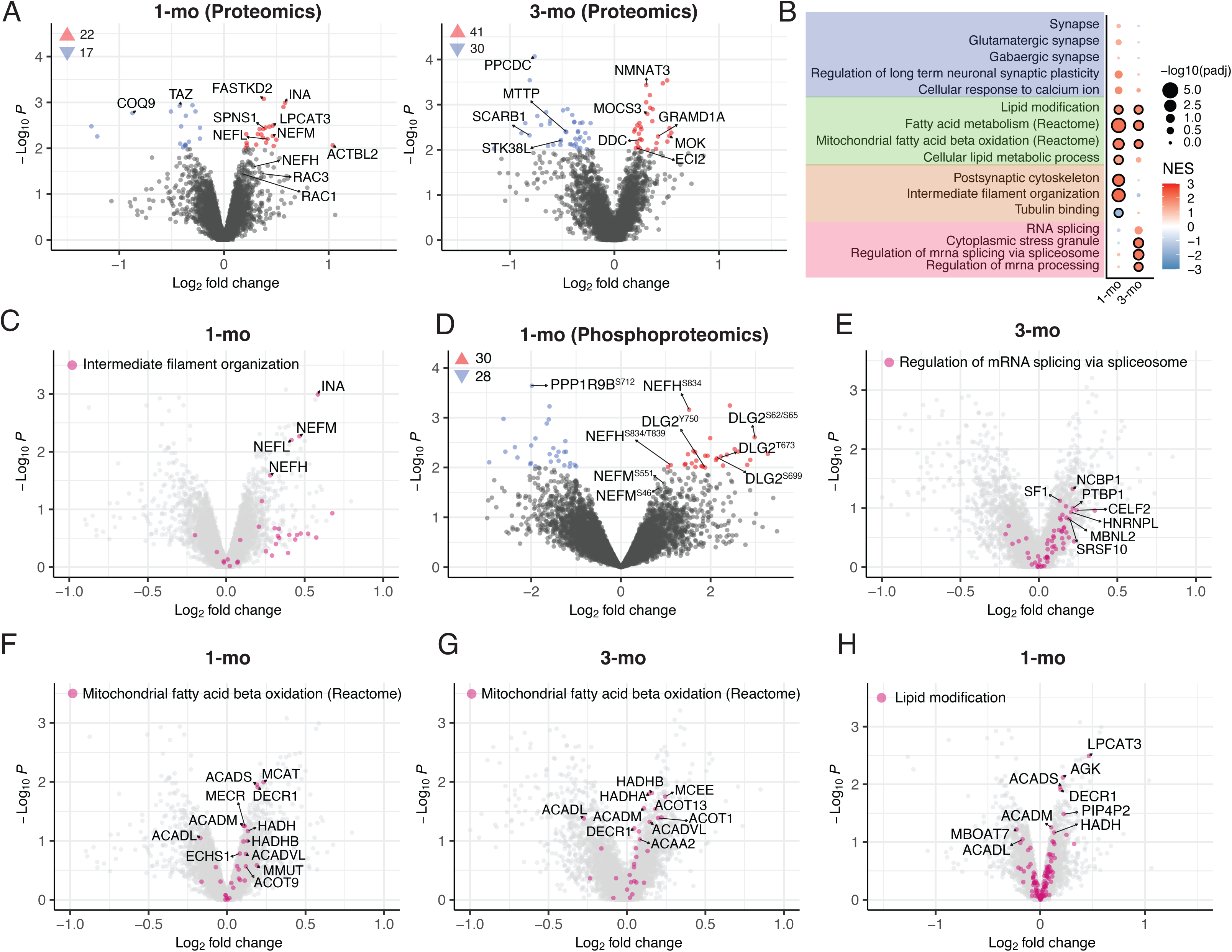
Alterations in the synaptic proteome of the *Zmym2*^+/-^ cortex (A) Volcano plots showing proteomic changes in synapses of 1-& 3-mo *Zmym2*^+/-^ mice. Red and blue dots represent DEPs (p < 0.01). (B) GSEA results of the synaptic proteome in 1-and 3-mo *Zmym2*^+/-^ mice. Reactome: gene list from the Reactome Pathway Database. (C, E-H) Volcano plots showing proteomic changes in indicated brain regions of *Zmym2*^+/-^ synapses, highlighting proteins within selected gene sets. (D) Volcano plot of phosphoproteomic changes in synapses of 1-mo *Zmym2*^+/-^ mice. Phosphoproteomic measurements were normalized to proteomic measurements from the same samples. Red and blue dots represent differentially phosphorylated proteins with specific phosphorylation sites (p < 0.01).

It is notable that GSEA of *Zmym2*^+/-^ synaptic proteomics showed little signal for synapse-or glutamatergic synapse-related gene sets (Fig. 4B), and the core components of the synapse (such as glutamate receptors and postsynaptic scaffolds, presynaptic vesicle release machinery) were relatively unaltered in abundance. On the other hand, GO terms related to neurofilaments— critical for neuronal structure—were prominently upregulated in the synapse proteome at 1-mo, with significant elevation of all three major neurofilament subunits (NEFL, NEFM, NEFH) and the associated protein INA (α-internexin) (Fig. 4B, C). This elevation in neurofilament subunits is intriguing, as increased NEFL levels are emerging as biomarkers for neurodegenerative disorders and perhaps mental illnesses (Khalil et al., 2018). In contrast, a number of proteins involved in tubulin binding, such as TUBGCP4, KIF3C, and MAP9 (Fig. 4B, Supplemental Fig. 7B), as well as proteasome subunits (Supplemental Fig. 7C, D), were downregulated.

The expression of neurofilament subunits at the RNA level was unchanged or decreased at 1-mo in the *Zmym2*^+/-^ brain (Supplemental Fig. 7E), implying that post-transcriptional mechanisms underlie the increased protein levels of neurofilaments. Phosphorylation of neurofilament subunits is known to play a critical role in stabilizing neurofilament structure and reducing their susceptibility to degradation (Goldstein et al., 1987; Pant, 1988). Through quantitative phosphoproteomics, we identified increased phosphorylation of neurofilament subunits, including NEFH and NEFM at multiple sites (Fig. 4D, Supplemental Table 5).

Phosphorylation states of synaptic scaffold proteins are also known to regulate synaptic functions; for example, synaptic strength is influenced by the phosphorylation of PSD-95 (Kim et al., 2007; Nelson et al., 2013). Our phosphoproteomics analysis revealed increased phosphorylation of DLG2 (PSD-93; Fig. 4D), a PSD-95-family protein whose phosphorylation enhances its interaction with synaptic components and regulates structural synaptic plasticity (Hossen et al., 2022; Parker et al., 2004). Together, these synaptic proteomics and phosphoproteomics findings suggest that proteins associated with structural integrity and scaffolding at the synapse are dysregulated in *Zmym2*^+/−^ mice at 1-mo.

At 3-mo, GO terms related to mRNA splicing/stress granules showed upregulation in synapstic proteomics of *Zmym2*^+/-^ mice (Fig. 4B, E, Supplemental table 6), a finding also in the synaptic proteomics of *Grin2a* mutant mice (Farsi et al., 2023). For instance, NCBP1, a protein implicated in pre-mRNA splicing (Izaurralde et al., 1994) and cognitive function (Gao et al., 2023), is increased in the synaptic fractions of both *Zmym2* (Fig. 4E) and *Grin2a* mutant mice (Farsi et al., 2023). The heightened presence of RNA processing and stress granule-associated proteins in synaptic fractions might suggest mislocalization of these RNA-binding proteins from the nucleus to the cytoplasm and synapse, mechanisms which are implicated in neurodegenerative conditions such as amyotrophic lateral sclerosis (ALS) and frontotemporal dementia (Luan et al., 2023).

A particularly prominent proteomics finding at both 1-and 3-mo was the upregulation of lipid metabolism pathways (Fig. 4B, F, G, H, Supplemental Table 6). Proteins involved in beta oxidation, a process that breaks down fatty acids to produce energy in mitochondria, were elevated at both developmental stages (Fig. 4B, F, G). Additionally, several proteins associated with lipid peroxidation were increased in abundance in 1-mo *Zmym2* mutant mice. For instance, LPCAT3 (lysophosphatidylcholine acyltransferase 3), an enzyme involved in lipid remodeling processes and a modulator of ferroptosis sensitivity (Cui et al., 2023; Reed et al., 2022), showed increased abundance (Fig. 4A, H). Similarly, DECR1 (2,4-dienoyl-CoA reductase 1), a rate-limiting enzyme in the oxidation of polyunsaturated fatty acids and a known regulator of ferroptosis (Blomme et al., 2020; Nassar et al., 2020), was also elevated (Fig. 4H). These findings suggest that dysregulated lipid metabolism, which can affect ferroptosis and synaptic function (Valles and Barrantes, 2022), is a prominent feature in the brains of *Zmym2* mutant mice.

### Metabolomic alterations in the cortex of *Zmym2* mutant mice

Given that metabolism-related GO terms were significantly enriched in our GSEA of proteomic data (Fig. 4B), we employed unbiased liquid chromatography-tandem mass spectrometry (LC-MS/MS) to investigate metabolomic changes in total lysates from the cerebral cortex of 1-mo *Zmym2*^+/-^ mice. Our profiling identified 829 annotated metabolites across 16 categories, with complex lipids, including glycerophospholipids, glycerolipids, and sphingolipids, constituting the largest group. Organic acids accounted for 12% (101 out of 829) of the metabolome, while nucleic acids contributed 6% (Supplemental Fig. 8A).

We identified 36 metabolites with significant changes (p < 0.05) in the *Zmym2*^+/-^ cortex (Fig. 5A, Supplemental Table 7). Most sphingomyelin (SM) species were decreased in abundance (e.g., SM 36:1;O2, SM 34:2;O2, SM 36:2;O2, SM 36:0;O2), whereas most ceramides (Cers) were increased in the *Zmym2*^+/-^ cortex (e.g., Cer 38:1;O2, Cer 36:1;O2) (Fig. 5B). SMs and Cers are enriched in lipid rafts (Bieberich, 2018) and in myelin (Yoo et al., 2020), both of which are important for synaptic function and neuronal communication (Hering et al., 2003; Zemmar et al., 2018).

**Figure 5:**
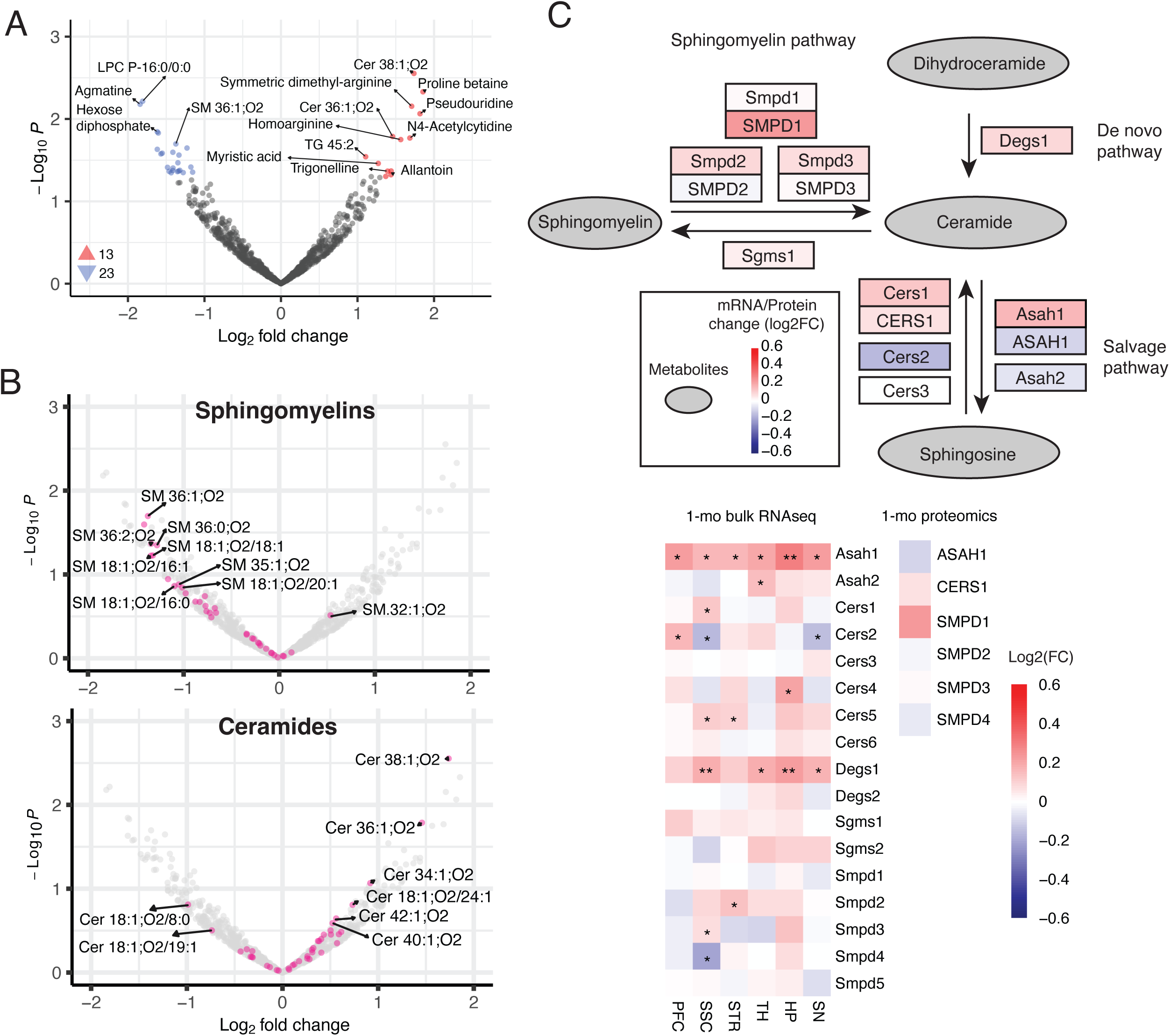
Metabolomic alterations in the *Zmym2*^+/-^ cortex (A) Volcano plot showing metabolomic changes in *Zmym2*^+/-^ mice, with red and blue dots indicating metabolites with p < 0.05. (B) Volcano plots highlighting metabolites related to selected categories in *Zmym2* mutants. (C) Simplified overview of the SM and Cer metabolism pathway. The color of squares uppercase gene names represents log2FC values from 1-mo proteomics data, while the color of squares containing lowercase gene names (except for the first letter) represents log2FC values from bulk RNA-seq data of the 1-mo SSC. The heatmap displays log2FC values of genes/proteins involved in SM and Cer metabolism, derived from bulk RNA-seq and proteomics analyses of 1-mo *Zmym2*^+/-^ mice. *: p < 0.05; **: padj < 0.05

Cer levels have been reported to be elevated in brain tissues from individuals with Alzheimer’s disease (AD) (Filippov et al., 2012) and SCZ (Schwarz et al., 2008). What might explain the increased levels of Cers in the *Zmym2* mutant mouse brain? As shown in Fig. 5C, we found that *Degs1*, which encodes delta(4)-desaturase—an enzyme that catalyzes the conversion of dihydroceramide to Cer—was significantly upregulated at the RNA level in multiple brain regions of 1-mo *Zmym2*^+/-^ mice (Fig. 5C). Additionally, *Smpd1*, which encodes an enzyme that catalyzes the hydrolysis of SMs into Cers, was increased at both RNA and protein levels, as was *Cers1*, which synthesizes Cers from sphingosine (Fig. 5C).

The ACSL4–LPCAT3–lipoxygenases (LOX) pathway plays a critical role in lipid peroxidation, which is a trigger for ferroptosis, a regulated form of cell death characterized by the accumulation of lipid peroxides (Gupta et al., 2023; Zhuo et al., 2024). In *Zmym2*^+/-^ mice, LPCAT3 protein was increased in synaptic fractions at 1-mo (Fig. 4A) and was also elevated across multiple brain regions at the RNA level (Supplemental Fig. 8B). Several genes encoding LOX enzymes, which oxidize polyunsaturated fatty acids to generate lipid hydroperoxides, including *Alox8* and *Alox12b*, were upregulated across the brain, particularly in the PFC and SSC (Supplemental Fig. 8B). This was accompanied by an overall upregulation of LOX-related pathways across most brain regions (Supplemental Fig. 8C). Notably, protein level of GPX4, a key enzyme that reduces lipid hydroperoxides to non-toxic lipid alcohols and counteracts ferroptosis (Seibt et al., 2019), was also upregulated (Supplemental Fig. 8B).

We further analyzed the levels of hydroxy derivatives from LOX-mediated lipid peroxidation, which are more stable than their hydroperoxide precursors (Dobrian et al., 2011) and serve as indicators of LOX activity (Pratico et al., 2004). Our untargeted metabolomics analysis revealed an increase in almost all LOX-derived metabolites, including hydroxyeicosatetraenoic acids (HETEs), hydroxyoctadecadienoic acids (HODEs), and hydroxydocosahexaenoic acids (HDoHEs), in *Zmym2*^+/-^ mutants, though no individual molecular change reached statistical significance (Supplemental Fig. 8B) (Ackermann et al., 2017; Kulkarni et al., 2021). Elevated levels of HETEs and HODEs have been reported in several neurological disorders (Shichiri, 2014), for example, increased levels of 12-and 15-HETE (Pratico et al., 2004; Yao et al., 2005) and total HODE (Yoshida et al., 2009) have been observed in AD patients. Together, our transcriptomic and metabolomic analyses suggest enhanced lipid peroxidation in *Zmym2*^+/-^ mice, which together with other changes in lipid metabolism, may contribute to the pathophysiology.

### EEG abnormalities in *Zmym2* mutant mice

Brain pathophysiology can be reflected in aberrant EEG patterns. Abnormal EEG features are commonly observed in individuals with SCZ and ASD (Kozhemiako et al., 2022; Nicotera et al., 2019) as well as in mouse models of these disorders (Aryal et al., 2024; Herzog et al., 2023; Martinez et al., 2024). To investigate the impact of *Zmym2* LoF on brain network activity, we performed 24-hour EEG recordings in *Zmym2*^+/-^ mutant mice, using electrodes placed over the frontal and parietal cortices. The duration of wake, REM, and NREM phases during the light and dark cycles was similar between *Zmym2*^+/-^ mice and their WT littermates at both 3-and 6-mo (data not shown).

We analyzed the power of brain oscillations across multiple frequency bands during NREM sleep, a phase less affected by movement-related EEG artifacts. *Zmym2*^+/-^ mutants exhibited significant genotype differences in the frontal and parietal cortices at 3-and 6-mo (relative oscillatory power across all major frequency bands of brain oscillation, Fig. 6A, B). Notably, delta oscillations, which have been reported to show increased power in SCZ (Maki-Marttunen et al., 2019; Sponheim et al., 1994), were significantly elevated in relative power in *Zmym2*^+/-^ mice (Fig. 6C, D). Conversely, theta oscillations, which are associated with memory processes (Khader et al., 2010; Larrain-Valenzuela et al., 2017), were significantly reduced in relative power in *Zmym2*^+/-^ mice (Fig. 6C, D), a pattern also observed in wake-phase EEG (Supplemental Fig. 9A).

**Figure 6:**
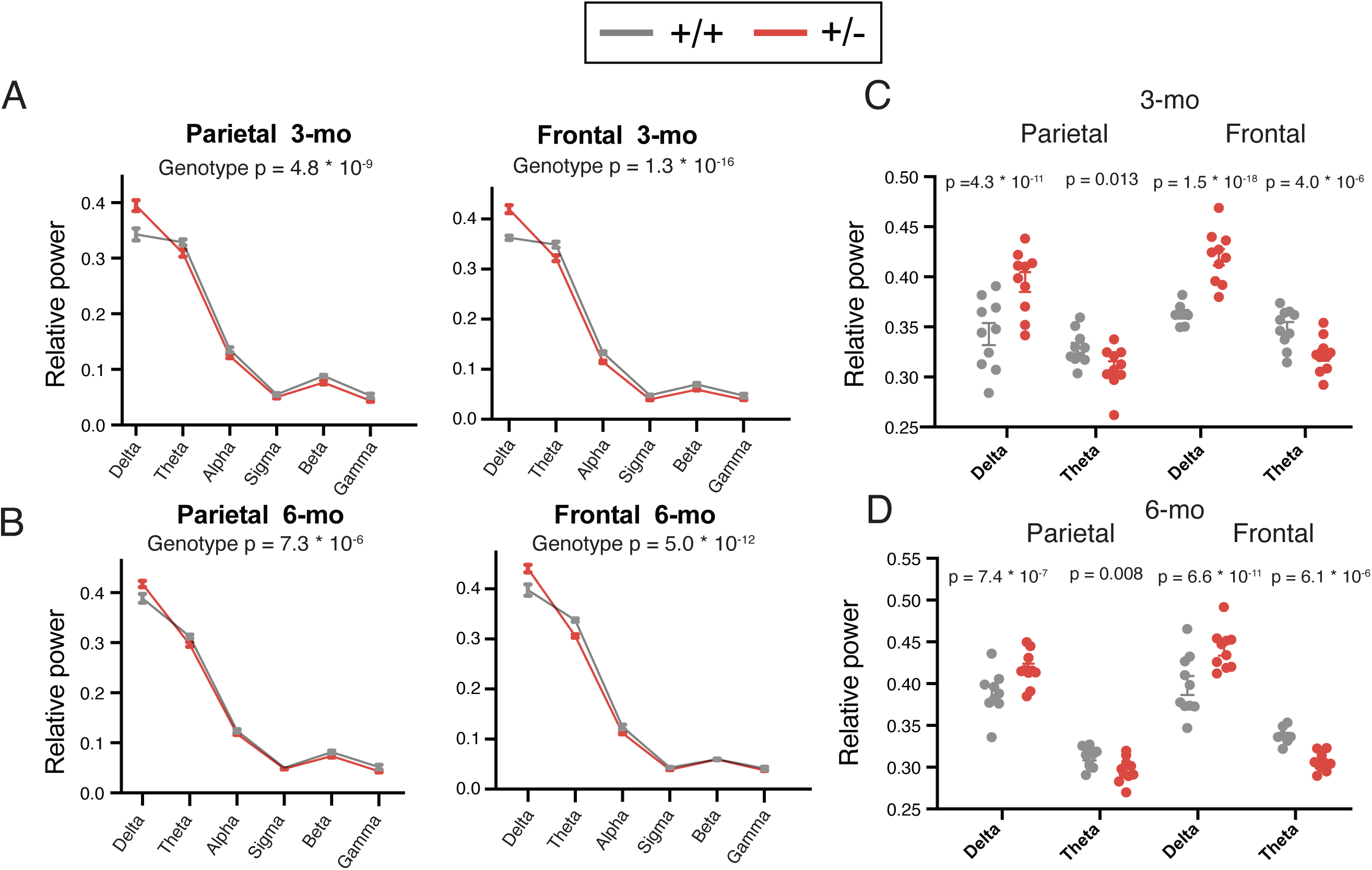
Abnormal EEG patterns in the *Zmym2*^+/-^ cortex (A, B) Relative NREM power spectra in the frontal and parietal cortex in 3-and 6-mo *Zmym2* mutants during NREM sleep. The p values in the plots were calculated using a linear mixed-effects model to test for genotype differences. (C, D) Relative delta and theta power in the frontal and parietal cortex during NREM sleep at 3-and 6-mo in *Zmym2*^+/-^ mice and WT controls. The p values were calculated by a linear mixed-effects model to test for genotype differences.

Additionally, we assessed sleep spindles, which are potentially linked to memory consolidation and are reduced on average in individuals with SCZ (Kozhemiako et al., 2024; Kozhemiako et al., 2022). However, no significant differences in sleep spindle density, amplitude, or duration were observed between genotypes (Supplemental Fig. 9B-D).

### Locomotor hyperactivity in *Zmym2* mutant mice

Amphetamine-induced hyperlocomotion in rodents is a commonly used behavioral model for psychosis (Kalivas and Stewart, 1991), and mouse mutants of *Grin2a* show locomotor hyperactivity (Herzog et al., 2023). In the open field test (OFT), 1-mo male *Zmym2*^+/-^ mice exhibited increased locomotor activity (∼28% increase in total distance traveled compared to WT; Fig. 7A-C), with no significant difference in center/margin ratio (Fig. 7C). The difference in *Zmym2*^+/-^ locomotor activity diminished with age and was not statistically significant at 3-mo (Fig. 7C). Female *Zmym2*^+/-^ mice showed no significant difference in locomotor activity and center/margin ratios in OFT (Supplemental Fig. 10A-C).

**Figure 7:**
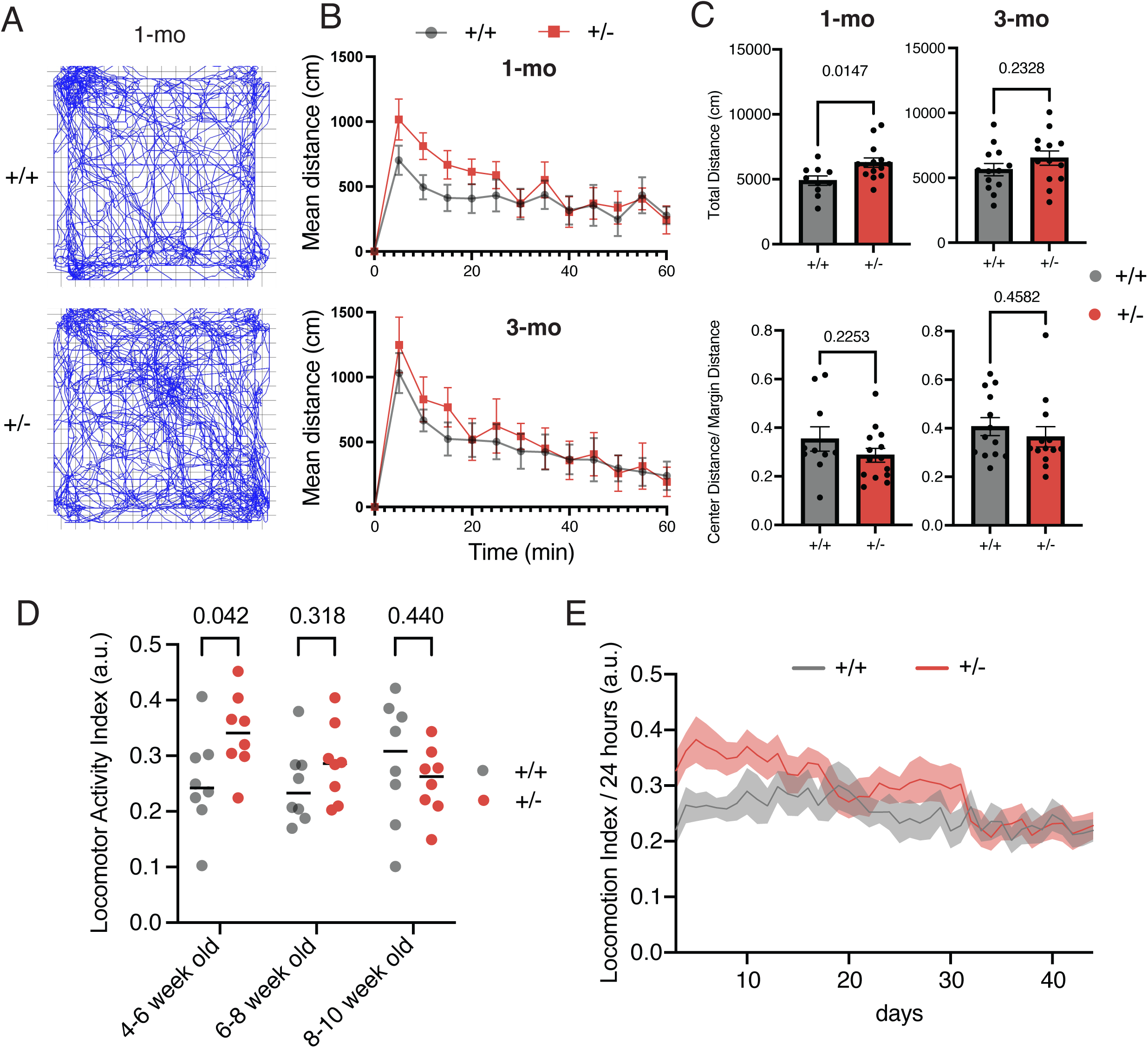
Locomotor behavior of *Zmym2*^+/-^ mice in the OFT and homecage monitoring analysis (A) Representative movement traces of 1-mo *Zmym2*^+/-^ mice during a 60-min OFT. (B) Mean distance traveled, binned into 5-minute intervals. (C) Total distance traveled and center/margin distance ratio for 1-mo (left) and 3-mo (right) mice. Error bars represent mean ± standard error. The p values from two-tailed t-tests are indicated in each plot. (D) Locomotor activity index across three developmental stages, with p values shown for each comparison (two-tailed t-test). (E) Mean locomotor activity index across monitoring during day 1 to day 45.

Using the Digital Ventilated Cage (DVC) system for home cage monitoring (Iannello, 2019), we recorded various behavioral metrics including locomotor activity beginning at 1-mo for 1.5-2 months. Consistent with the OFT results at 1-mo, *Zmym2*^+/-^ mice exhibited higher locomotor activity in their home cages at 4-6 weeks of age; however, this hyperactivity was no longer statistically significant at 6-8 weeks or 8-10 weeks of age (Fig. 7D, E). In female mutant mice, no significant differences in locomotor activity were observed at 4 to 10 weeks of age (Supplemental Fig. 10D, E).

## DISCUSSION

In this study, we investigated the effects of *Zmym2* heterozygous LoF (mimicking the human disease-linked genetic variant, which is linked to SCZ, ASD, and DD/ID) on the mouse brain using transcriptomics, proteomics, metabolomics, EEG, and behavioral approaches. Consistent with ZMYM2’s role as an integral component of several transcriptional regulatory complexes, *Zmym2* heterozygous LoF resulted in widespread disturbances of gene expression, including de-repression of germline genes and dysregulated expression of epigenetic regulators such as HDAC1. Although ZMYM2 primarily functions in the nucleus, its deficiency impacted neuronal activity, synapses and synaptic function, as evidenced by downregulation of ARGs and synapse-related genes in neurons and altered protein composition of synapses.

Epigenetic mechanisms play a crucial role in regulating gene expression and are often affected in individuals with SCZ (Smigielski et al., 2020) and ASD (Waye and Cheng, 2018). It is therefore interesting that ZMYM2, a protein associated with several epigenetic complexes, has been identified as a high-impact LoF risk gene for SCZ. Among the 32 risk genes identified in the SCHEMA study (Singh et al., 2022), approximately 20% (6 out of 32; *SETD1A*, *KDM6B*, *ASH1L*, *HIST1H1E*, *STAG1*, and *ZMYM2*) are implicated in epigenetic regulation (Farsi and Sheng, 2023). Similarly, many high-confidence ASD risk genes, including *ADNP*, *CHD8*, *POGZ*, and *ARID1B*, are known to modulate epigenetic functions (Satterstrom et al., 2020), suggesting that disruptions in epigenetic regulation may contribute to the pathophysiology of both SCZ and ASD. Consistent with this notion, our findings indicate that in *Zmym2* mutants, (i) germline genes are aberrantly de-repressed across multiple brain regions (Fig. 1C), and (ii) pathways related to histone modification and chromatin remodeling are broadly downregulated (Fig. 3A).

In keeping with *ZMYM2*’s central role in regulating gene expression, *Zmym2*^+/-^ mice exhibit widespread alterations in gene expression across brain regions (Fig. 1A, 2A). One of the most striking gene expression changes is the upregulation of HDAC1 in multiple brain regions, particularly in the STR (Fig. 3C, D). HDAC1, a member of the class I HDAC family, has been implicated in both ASD and SCZ. For instance, inhibition of class I HDACs has been shown to alleviate social deficits and enhance the expression of synaptic and actin-regulator genes in SHANK3-deficient mice (Qin et al., 2018). Elevated HDAC1 levels and reduced expression of HDAC1-binding genes, such as *Gad1* and *Pvalb*, have been observed in the PFC of SCZ patients (Bahari-Javan et al., 2017). Further investigation into the dysregulation of epigenetic regulators, particularly HDAC1, may provide insights into the pathological mechanisms underlying ZMYM2 deficiency.

ARGs, which can be measured as transcriptomic markers of neuronal activity, were reduced in several brain regions of *Zmym2* mutants. Transcriptomic analysis pointed to PFC hypoactivity (Fig. 2C), which is consistent with findings in individuals with SCZ (Minzenberg et al., 2009) and in mouse models with deficiencies in SCHEMA genes, such as *Grin2a*^+/-^, *Gria3*^-/y^, and *Srrm2*^+/-^ mice (Aryal et al., 2024; Farsi et al., 2023; Huang et al., 2024). In both bulk and snRNA-seq analyses (Fig. 1C and 2C), the STR exhibited downregulation of ARGs, suggesting hypoactivity. This observation is different from the STR hyperactivity that was reported in individuals with SCZ (Li et al., 2020) and in *Grin2a* mouse mutants (Farsi et al., 2023). However, it aligns with findings in *Srrm2*^+/-^ mutant mice, which carry LoF mutations in another gene associated with SCZ and NDD (Aryal et al., 2024). More direct assessments of neuronal activity, such as neuroimaging or electrophysiological recordings, will be essential for further elucidating the pathophysiology of *Zmym2* mutant mice.

There is evidence of elevated Cer levels in the plasma and brain tissue of individuals with neuropsychiatric and neurodegenerative disorders (Bernal-Vega et al., 2023). For example, genomic and lipidomic studies in individuals with SCZ have demonstrated that (i) a genome-wide association study identified three SNPs in *SMPD3*, which encodes an enzyme involved in Cer production, that showed a significant association with SCZ (Chestnykh et al., 2025), and (ii) Cer levels were increased in brain samples from individuals with SCZ (Esaki et al., 2020; Schwarz et al., 2008). Furthermore, elevated levels of lipid peroxidation markers in plasma and cerebrospinal fluid suggest increased lipid peroxidation in SCZ (Mahadik et al., 1998; McCreadie et al., 1995). In *Zmym2*^+/-^ mice, we observed an overall increase in Cer levels (Fig. 5B) as well as LOX-derived metabolites (Supplemental Fig. 8B). A potential link between Cer accumulation and lipid peroxidation is increased oxidative stress, which may contribute to the pathology of neuropsychiatric and neurodegenerative disorders (Adibhatla and Hatcher, 2010; Jazvinscak Jembrek et al., 2015). Follow-up studies investigating oxidative stress could provide insights into the molecular basis of lipid composition alterations in the *Zmym2*^+/-^ brain.

Overall, this comprehensive, unbiased-omics study provides new insights into the molecular and neurobiological consequences of *ZMYM2* mutations, as well as potential mechanisms underlying these effects, thereby advancing our understanding of SCZ and NDD pathophysiology.

## Acknowledgements

We thank Kris Dickson for valuable input for this manuscript; Sean K. Simmons and Joshua Z. Levin for their advice on transcriptomic analysis; Magdalena Sevilla-Gonzalez for sharing her pipeline and input on bioinformatic analysis of metabolomics.

## Author contributions

W.-C.H. and M.S. conceived and designed the experiments. The manuscript was written by W.-C.H. and M.S., with input from all authors. W.-C.H. performed sample and library preparations for bulk and snRNA-seq, conducted data analysis and visualization, and carried out behavioral tests.

K.P.M. processed the transcriptome data, including quality control, alignment, cell type clustering, annotation, and differential expression analysis. S.A. purified synapse fractions for proteomics. A.V.-T. performed LC-MS/MS for synaptic proteomics under the guidance of H.K. and S.A.C. C.G. performed immunoblots. E.M., C.D., L.D., A.D., and L.I. prepared samples and performed LC-MS/MS for metabolomics under the guidance of C.B.C. Z.F. assisted with data analysis on transcriptomics. X.-M.L. assisted with data analysis on metabolomics. N.G., A.S.A., and Y.W. performed data acquisition and analysis of EEG under the guidance of J.Q.P. B.J.S. set up the home cage monitoring system and assisted with testing and data analysis.

## FIGURE LEGENDS

**Supplemental Fig. 1: Gross metrics of *Zmym2*^+/-^ mutant mice**

(A) Number of mice with each genotype from *Zmym2*^+/-^ x *Zmym2*^+/-^ breeding. (B) Relative *Zmym2* mRNA levels measured by qRT-PCR, normalized to *Gapdh* mRNA. Error bars denote mean ± standard error. The p value from a two-tailed t-test is shown. (C) Western blot analysis of ZMYM2 and ACTB in cortical lysates from 1-mo *Zmym2* animals. (D-F) Bar charts representing body weight, brain weight, and body length in mice. Error bars represent mean ± standard error; the p value for each chart is indicated (two-tailed t-test).

**Supplemental Fig. 2: Correlated transcriptomic changes across different brain regions in *Zmym2*^+/-^ mice**

(A) Venn diagram showing overlapping DEGs across brain regions at three developmental stages. Numbers indicate the number of DEGs. (B) Correlation of RNA-seq results across brain regions, with Spearman’s correlation was performed on log2FC values for each region pair. Numbers in squares represent Spearman’s correlation coefficient. (C) Comparison of correlations between *Zmym2* and *Grin2a* transcriptomic profiles, with each dot representing a correlation coefficient for a brain region pair. (D) Heatmap showing log2FC values of three active LINE-1 elements in bulk RNA-seq data from the indicated brain regions in *Zmym2*^+/-^ mice. *: p < 0.05; **: padj < 0.05.

**Supplemental Fig. 3: GSEA results from bulk RNA-seq in *Zmym2*^+/-^ mutant mice with different gene sets**

(A, D) GSEA of bulk RNA-seq data using SynGO (A) and published disease gene lists (D), in *Zmym2*^+/-^ mice from indicated brain regions and ages. PTSD: post-traumatic stress disorder, ADHD: attention deficit hyperactivity disorder, MSA: multiple system atrophy, and ALS: amyotrophic lateral sclerosis (B, C) Volcano plots of transcriptomic changes in the indicated brain regions of *Zmym2* mutants, highlighting genes within the specified gene sets. Dot plots showing transcriptomic changes between 1-and 3-mo *Zmym2* mutants, with Spearman’s correlation coefficient (r) for log2FC between these ages.

**Supplemental Fig. 4: Transcriptomic changes in female *Zmym2*^+/-^ mutant mice**

(A) Number of DEGs in the indicated brain region in 1-mo female *Zmym2^+/-^*mutants. Circle size represents number of DEGs (padj < 0.05), while color represents expression direction. (B) Expression changes of genes in bulk RNA-seq from female *Zmym2*^+/-^ mice in indicated brain regions, with circle size indicating statistical significance and log2FC by color scale. Circles with black outlines indicate padj < 0.05. (C) GSEA of bulk RNA-seq in 1-mo female and male *Zmym2*^+/-^ from indicated brain regions. Statistical significance is shown by circle size, and NES by color scale. Circles with black outlines indicate padj < 0.05. Male results, originally presented in Figure 1C, are re-displayed here for comparison. (D) Volcano plot of transcriptomic changes in 1-mo PFC of *Zmym2*^+/-^ female mice, highlighting genes in the “meiosis I cell cycle” gene set.

**Supplemental Fig. 5: Identification of diverse cell types by snRNA-seq analysis in *Zmym2*^+/-^ mouse brain regions**

(A) Fraction of each identified cell type in the indicated brain regions and developmental stages, with each column representing the average value per genotype (n = 4-6 animals per genotype). (B) GSEA of snRNA-seq in *Zmym2*^+/-^ mice across the indicated brain regions and developmental stages. (C) Volcano plots of transcriptomic changes in the PFC of *Zmym2* mutants, highlighting genes in the “Steroid biosynthetic process” gene set.

**Supplemental Fig. 6: Comparison of transcriptomic changes between *Zmym2*, *Adnp*, and *Pogz* mutant mice**

(A) Overlap of DEGs across six brain regions or (B) adult HP across three mouse lines. Numbers indicate the number of DEGs. (C) Heatmap of log2FC values from bulk RNA-seq data from indicated brain regions in *Adnp*, *Pogz*, and *Zmym2* mutant mice. **: padj or FDR < 0.05; *: p < 0.05. The genes identified in all three mouse lines were used for comparison. (D) Heatmap of NES values for overlapping GO terms from adult HP across three mouse lines. (E) GSEA results for selected GO terms in indicated brain regions of *Adnp*, *Pogz*, and *Zmym2* mutants, with circle size indicating statistical significance and NES is indicated by the color scale. Circles with black outlines indicate padj < 0.05.

**Supplemental Fig. 7: Alteration in synaptic proteome of the *Zmym2*^+/-^ cortex**

(A) Dot plot showing proteomic changes in 1-and 3-mo *Zmym2* mutants. Spearman’s correlation coefficient (r) represents the correlation of log2FC values of DEPs between 1-and 3-mo. (B, D) Volcano plots of proteomic changes in *Zmym2* mutants, highlighting proteins within specific gene sets. (C) GSEA of synaptic proteome from indicated developmental stages, with circle size indicating statistical significance and NES indicated by color scale. Circles with black outlines indicate padj < 0.05. Reactome: gene list from the Reactome Pathway Database; WP: gene list from the WikiPathways Database (E) Heatmap showing log2FC values of neurofilament subunits from bulk RNA-seq data in the indicated brain regions of *Zmym2*^+/-^ mice. *: p < 0.05; **: padj < 0.05

**Supplemental Fig. 8: Metabolomic alterations in the *Zmym2*^+/-^ cortex**

(A) Chemical composition of the brain metabolome, categorized by UCSD Metabolomics Workbench classifications to illustrate the diversity among annotated metabolites. (B) Simplified overview of the ACSL4-LPCAT3-LOX pathway. The color of squares containing uppercase gene names represents log2FC values from 1-mo proteomics data, while the color of squares containing lowercase gene names (except for the first letter) represents log2FC values from bulk RNA-seq data of the 1-mo SSC. The heatmap displays log2FC values of genes/proteins related to the ACSL4-LPCAT3-LOX pathway, derived from bulk RNA-seq and proteomics analyses of 1-mo *Zmym2*^+/-^ mice. *: p < 0.05; **: padj < 0.05. PUFA: polyunsaturated fatty acid; PUFA-CoA: coenzyme A-activated polyunsaturated fatty acid; PL-PUFA: PUFA-containing phospholipids; PUFA-PL-OOH: PUFA phospholipid hydroperoxides; PL-PUFA-OH: PUFA phospholipid alcohols; NE-PUFA-OOH: non-esterified PUFA hydroperoxides; NE-PUFA-OH: non-esterified hydroxy-PUFAs. Volcano plots highlighting metabolites associated with selected categories in *Zmym2* mutants. (C) GSEA results from bulk RNA-seq analysis in *Zmym2*^+/-^ mice, with circle size indicating statistical significance and NES indicated by the color scale. Circles with black outlines indicate padj < 0.05

**Supplemental Fig. 9: Abnormal EEG patterns in the *Zmym2*^+/-^ cortex**

(A) Relative power spectra in the frontal and parietal cortex of 3-and 6-mo *Zmym2* mutants during the wake phase. The p values were calculated using a linear mixed-effects model to test for genotype differences. Asterisks indicate p < 0.05 in the specified frequency bands. (B-D) Density, amplitude, and duration of 9 Hz, 11 Hz, 13 Hz, and 15 Hz spindles during NREM sleep. Error bars represent mean ± standard error. The p values in each plot were calculated using a linear mixed-effects model to test for genotype differences.

**Supplemental Fig. 10: Locomotor behavior of *Zmym2*^+/-^ female mice in the OFT and homecage monitoring analysis**

(A) Representative movement traces in 1-mo *Zmym2*^+/-^ female mice during a 60-min OFT. (B) Mean distance traveled, binned into 5-minute intervals. (C) Total distance traveled and center/margin distance ratio in 1-mo (left) and 3-mo (right) *Zmym2*^+/-^ female mice. Error bars represent mean ± standard error. The p values for each plot were from two-tailed t-tests. (D) Locomotor activity index across three developmental stages, with p values for each comparison (two-tailed t-test). (E) Mean locomotor activity index from monitoring from day 1 to day 45.

## METHODS

### EXPERIMENTAL MODEL

*Zmym2*^+/-^ mice (C57BL/6NJ-Zmym2^em1(IMPC)J/Mmjax^; #051239-JAX) were originally obtained from the Jackson Laboratory (JAX). They were generated by the Knockout Mouse Phenotyping Program (KOMP) using CRISPR to delete approximately 300 base pairs in exon 4. *Zmym2* heterozygous mutant (*Zmym2*^+/-^) mice were initially bred by crossing *Zmym2*^+/-^ males and females. After approximately 10 litters without obtaining any *Zmym2*^-/-^ offspring, we adjusted our breeding strategy by crossing *Zmym2*^+/-^ males with C57BL/6NJ females (#005304), as recommended by JAX for control mice. The mice were maintained on a 12-hour light/dark cycle with *ad libitum* access to food and water. All procedures were approved by the Broad Institute Institutional Animal Care and Use Committee (IACUC). *Zmym2* mutant mice and their WT littermates at 1-, 3-, 5-mo were used in this study.

## METHOD DETAILS

### Brain perfusion and dissection

The detailed procedure of brain perfusion and dissection was described previously (Farsi et al., 2023). In brief, mice were anesthetized with isoflurane, and transcardial perfusions were performed with ice-cold HBSS (Life Technologies) to remove blood from the brain, which was then immediately frozen in liquid nitrogen vapor and stored at –80°C. Brain dissection was carried out in a cryostat, with all tools precooled to –20°C. The cerebellum was removed first, and then the medial PFC, dorsal HP, TH, SSC, and dorsal STR, and SN, which were meticulously dissected using a biopsy punch and ophthalmic microscalpel in a cryostat (Leica). The brain regions were identified using the Allen Brain Atlas, and each excised tissue was stored at –80°C in 1.5mL tubes.

### Sample and library preparation for transcriptomic analysis

We followed previously published protocols to prepare samples and libraries (Farsi et al., 2023). For bulk RNA sequencing, RNA was extracted using the RNeasy Mini Kit (Qiagen), and its concentration and integrity were assessed using a NanoDrop Spectrophotometer and an Agilent 2100 Bioanalyzer with RNA PICO chips, ensuring all samples for sequencing had RNA integrity numbers (RINs) greater than 7 (n ≥ 5 for each brain region of each genotype). The purified RNA was stored at-80°C until library preparation. Libraries were prepared using the TruSeq Stranded mRNA kit from 200 ng of total RNA per sample, with the resulting cDNA libraries quantified using High Sensitivity DNA chips on an Agilent 2100 Bioanalyzer. A 10 nM normalized library pool was then sequenced on a NovaSeq S2 (Illumina) with a read length of 50 bases.

For snRNA-seq library preparation, we followed a published protocol for gentle, detergent-based dissociation to extract nuclei, which is available on protocols.io (www.protocols.io/view/frozen-tissue-nuclei-extraction-bbseinbe). Nuclei were collected and enriched using a Sony SH800 sorter at the Flow Cytometry Core facility of the Broad Institute. RNA from the extracted nuclei was captured using the Chromium Kit v3.1 (10x Genomics), and library preparation followed the manufacturer’s guidelines. A 10 nM normalized library was pooled, and sequencing was conducted on a NovaSeq S2 (Illumina) with a customized paired-end, dual indexing format, as recommended by 10x Genomics. n ≥ 4 for each genotype.

### Quantitative Real-time PCR

RNA was extracted using the RNeasy Mini Kit, and reverse transcription was performed with the iScript kit (Bio-Rad) according to the manufacturer’s protocol. Real-time PCR was conducted using PowerTrack SYBR Green Master Mix (Thermo Fisher Scientific) on a BioRad CFX384 Real-Time PCR Detection System. The primers used were: (1) *Zmym2*: F: ATGACAGGCTCAGCACCAC; R: CACAGGGACAGGAATAGGC, and (2) *Gapdh*: F: GGCATTGCTCTCAATGACAA; R: CCCTGTTGCTGTAGCCGTAT. A two-tailed t-test was used for quantification comparisons.

### Purification of synaptic fraction for MS and quantitative MS analysis

The methods for this study were adapted from our previous papers (Aryal et al., 2023; Aryal et al., 2024; Farsi et al., 2023). Briefly, 1-mo mice were sacrificed using CO_2_ anesthesia, and their cortices were quickly dissected and flash-frozen in liquid nitrogen. For 3-mo, mice were perfused with HBSS and frozen (as described for RNA-seq), after which the cortices were dissected out in a cryostat. The cortical tissue was thawed, dounce-homogenized in ice-cold homogenization buffer (5 mM HEPES pH 7.4, 1 mM MgCl2, 0.5 mM CaCl2, supplemented with phosphatase and protease inhibitors), and centrifuged for 10 minutes at 1,400 g (4°C). The supernatant was further centrifuged at 13,800 g for 10 minutes (4°C), and the resulting pellet was resuspended in 0.32 M Sucrose, 6mM Tris-HCl (pH 7.5) and layered gently on a 0.85 M, 1 M, 1.2 M discontinuous sucrose gradient (in 6 mM Tris-HCl pH 8.0) and ultracentrifuged at 82,500 g for 2 hours (4°C). The band sedimenting at the 1 M and 1.2 M sucrose interface (the synaptosome fraction) was collected, treated with an equal volume of ice-cold 1% Triton X-100 (in 6 mM Tris-HCl pH 8.0), and incubated on ice for 15 minutes. The mixture was ultracentrifuged at 32,800 g for 20 minutes (4°C) to obtain the synapse fraction (postsynaptic density) pellet, which was solubilized and resuspended in 1% SDS. A small aliquot was used to measure the protein concentration of this fraction using the BCA micro assay (Pierce), and the remaining sample was stored at-80°C until being processed for mass spectrometry. Eighteen synaptic fractions from WT and *Zmym*2^+/-^ mouse cortices were analyzed using tandem mass tag (TMT) isobaric labeling for quantification (1-mo WT n = 4; 3-mo WT n = 4; 1-mo *Zmym2*^+/-^ n = 5; 3-mo *Zmym2*^+/-^ n = 5).

The proteins in synapse fraction samples (in 1% SDS) were reduced using 5 mM dithiothreitol and alkylated using 10 mM iodoacetamide at room temperature. The denatured, reduced, alkylated protein samples were then processed using S-Trap sample processing technology (Protifi) following manufacturer’s instructions. The proteins were bound to the S-Trap column via centrifugation and contaminants/detergents were washed away. Sequential digestion steps were then performed on column using 1:20 enzyme to substrate ratio of Lys-C for 2 hours and Trypsin overnight at room temperature.

Following digestion and desalting, 9 μg of each sample was labeled with TMT18 reagent; the TMT18 plex was constructed by randomly assigning the samples from each group to channels within the plex. After verifying successful labeling of more than 95% label incorporation, reactions were quenched using 5% hydroxylamine and pooled. The TMT18 labeled peptides were desalted on a 50 mg tC18 SepPak cartridge and fractionated on fractionated by high pH reversed-phase chromatography on a 2.1 mm x 250 mm Zorbax 300 extend-c18 column (Agilent). One-minute fractions were collected during the entire elution and fractions were concatenated into 12 fractions for LC-MS/MS analysis. Five percent of each fraction was removed for proteome analysis. The remaining portions of each of the 12 fractions were further concatenated to 6 fractions and dried down for phosphopeptide enrichment using immobilized metal affinity chromatography (IMAC).

One microgram of each proteome fraction was analyzed on a Exploris 480 QE mass spectrometer (Thermo Fisher Scientific) coupled to a Vanquish Neo LC system (Thermo Fisher Scientific). Samples were separated using 0.1% Formic acid / Water as buffer A and 0.1% Formic acid / Acetonitrile as buffer B on a 27 cm 75 um ID Picofrit column packed in-house with Reprosil C18-AQ 1.9 mm beads (Dr Maisch GmbH) with a 90 min gradient consisting of 1.8-5.4% B in 1 min, 5.4-27% B for 84 min, 27-54% B in 9 min, 54-81% B for 1 min followed by a hold at 81% B for 5 min. The MS method consisted of a full MS scan at 60,000 resolution and a normalized AGC target of 300% and maximum injection time of 10 ms from 350-1800 m/z followed by MS2 scans collected at 45,000 resolution with a normalized AGC target of 30% with a maximum injection time of 105 ms and a dynamic exclusion of 15 seconds. Precursor fit filter was used with a threshold set to 50% and window of 1.2 m/z. The isolation window used for MS2 acquisition was 0.7 m/z and 20 most abundant precursor ions were fragmented with a normalized collision energy (NCE) of 34 optimized for TMT18 data collection.

The data was analyzed using Spectrum Mill MS Proteomics Software (Broad Institute) with a mouse database from Uniprot.org downloaded on 04/07/2021 containing 55734 entries. Proteins identified by one or more peptides were selected for further analysis. Search parameters included: ESI Q Exactive HCD v4 35 scoring parent and fragment mass tolerance of 20 ppm, 40% minimum matched peak intensity, trypsin allow P enzyme specificity with up to four missed cleavages and calculate reversed database scores enabled. Fixed modifications were carbamidomethylation at cysteine. TMT labeling was required at lysine, but peptide N termini could be labeled or unlabeled. Allowed variable modifications were included protein N-terminal acetylation, oxidized methionine pyroglutamic acid, and pyro carbamidomethyl cysteine, as well as phosphorylated serine, threonine, and tyrosine for the phosphoproteome datasets. Protein quantification was achieved by taking the ratio of TMT reporter ions for each sample over the TMT reporter ion for the median of all channels. TMT18 reporter ion intensities were corrected for isotopic impurities in the Spectrum Mill protein/peptide summary module using the afRICA correction method which implements determinant calculations according to Cramer’s Rule and correction factors obtained from the reagent manufacturer’s certificate of analysis (https://www.thermofisher.com/order/catalog/product/90406) for lot numbers XJ346678 and XJ346678. After performing median-MAD normalization, a moderated two-sample t-test was applied to the datasets to compare WT and *Zmym2^+/-^* sample groups at 1 and 3-mo. A comprehensive list of DEPs for 1 and 3-mo *Zmym2*^+/-^ synapse fraction samples is provided in Supplemental Table 5.

### Western blotting

Western blotting was performed to quantify target protein levels. Total protein lysates were prepared by homogenizing mouse brain tissues in RIPA buffer (Thermo Fisher Scientific) and boiling the samples at 98°C for 7 minutes in 1X reducing SDS (Laemmli) buffer. Proteins were separated on 3–8% NuPAGE Bis-Tris gradient gels (Invitrogen) and transferred onto nitrocellulose membranes (BioRad). Membranes were blocked in EveryBlot Blocking Buffer (BioRad) for 60 minutes at room temperature, followed by overnight incubation at 4°C with primary antibodies (ZMYM2: Abcam (ab106624), HDAC1: Proteintech (10197-1-AP), β-Actin: Sigma (A3854), GAPDH: Proteintech (HRP-60004)). After four 5-minute washes, membranes were incubated with HRP-conjugated secondary antibodies for 1 hour at room temperature, washed again, and imaged using the ChemiDoc XRS+ system (BioRad). Data analysis was performed using ImageLab Software (BioRad), and a two-tailed t-test was used for protein quantification comparisons.

### Metabolomic analysis

Metabolomics on brain tissue was performed as liquid chromatography mass spectrometry (LC-MS) methods which measure both polar and non-polar metabolites. Brain tissues (n = 5 per genotype) were homogenized using a Qiagen TissueLyzer II in water at 1:4 (w:v) (TissueLyser II; Qiagen) and the aqueous homogenate was subjected to protein precipitation. Samples were prepared for each method using extraction procedures that are matched for use with the chromatography conditions. Data were acquired using LC-MS systems comprised of Nexera X2 U-HPLC systems (Shimadzu Scientific Instruments) coupled to Q Exactive/Exactive Plus orbitrap mass spectrometers (Thermo Fisher Scientific). The method details are summarized below.

### LC-MS method for HILIC-pos (positive ion mode MS analyses of polar metabolites)

LC-MS samples were prepared from media (10 μL) by protein precipitation with the addition of nine volumes of 74.9:24.9:0.2 v/v/v acetonitrile/methanol/formic acid containing stable isotope-labeled internal standards (valine-d8, Isotec; and phenylalanine-d8, Cambridge Isotope Laboratories). The samples were centrifuged (10 min, 10,000g, 4°C), and the supernatants injected directly onto a 150 × 2-mm Atlantis HILIC column (Waters). The column was eluted isocratically at a flow rate of 250 μL/min with 5% mobile phase A (10 mM ammonium formate and 0.1% formic acid in water) for 1 min followed by a linear gradient to 40% mobile phase B (acetonitrile with 0.1% formic acid) over 10 min. MS analyses were carried out using electrospray ionization in the positive ion mode using full scan analysis over m/z 70–800 at 70,000 resolution and 3-Hz data acquisition rate. Additional MS settings are: ion spray voltage, 3.5 kV; capillary temperature, 350°C; probe heater temperature, 300°C; sheath gas, 40; auxiliary gas, 15; and S-lens RF level 40.

### LC-MS method for HILIC-neg (negative ion mode MS analysis of polar metabolites)

LC-MS samples were prepared from media (30 μL) by protein precipitation with the addition of four volumes of 80% methanol containing inosine-15N4, thymine-d4 and glycocholate-d4 internal standards (Cambridge Isotope Laboratories). The samples were centrifuged (10 min, 10,000g, 4°C) and the supernatants were injected directly onto a 150 × 2.0-mm Luna NH2 column (Phenomenex). The column was eluted at a flow rate of 400 μL/min with initial conditions of 10% mobile phase A (20 mM ammonium acetate and 20 mM ammonium hydroxide in water) and 90% mobile phase B (10 mM ammonium hydroxide in 75:25 v/v acetonitrile/methanol) followed by a 10-min linear gradient to 100% mobile phase A. MS analyses were carried out using electrospray ionization in the negative ion mode using full scan analysis over m/z 60–750 at 70,000 resolution and 3 Hz data acquisition rate. Additional MS settings are: ion spray voltage, −3.0 kV; capillary temperature, 350°C; probe heater temperature, 325°C; sheath gas, 55; auxiliary gas, 10; and S-lens RF level 40.

### LC-MS method for C18-neg (negative ion mode analysis of metabolites of intermediate polarity; for example, bile acids and free fatty acids)

Media homogenates (30 μL) were extracted using 90 μL methanol containing PGE2-d4 as an internal standard (Cayman Chemical Co.) and centrifuged (10 min, 15,000g, 4°C). The supernatants (10 μL) were injected onto a Acquity HSS T3 C18 2.1 x 150mm (1.8um; Waters). The column was eluted isocratically at a flow rate of 450 μL/min with 20% mobile phase A (0.01% formic acid in water) for 3 min followed by a linear gradient to 100% mobile phase B (0.01% acetic acid in acetonitrile) over 12 min. MS analyses were carried out using electrospray ionization in the negative ion mode using full scan analysis over m/z 70–850 at 70,000 resolution and 3 Hz data acquisition rate. Additional MS settings are: ion spray voltage, −3.5 kV; capillary temperature, 320°C; probe heater temperature, 300°C; sheath gas, 45; auxiliary gas, 10; and S-lens RF level 60.

### LC-MS method for C8-pos

Lipids (polar and nonpolar) were extracted from media (10 μL) using 190 μL isopropanol containing 1-dodecanoyl-2-tridecanoyl-*sn*-glycero-3-phosphocholine as an internal standard (Avanti Polar Lipids; Alabaster, AL). After centrifugation (10 min, 10,000g, ambient temperature), supernatants (10 μL) were injected directly onto a 100 × 2.1-mm ACQUITY BEH C8 column (1.7 μm; Waters). The column was eluted at a flow rate of 450 μL/min isocratically for 1 min at 80% mobile phase A (95:5:0.1 v/v/vl 10 mM ammonium acetate/methanol/acetic acid), followed by a linear gradient to 80% mobile phase B (99.9:0.1 v/v methanol/acetic acid) over 2 min, a linear gradient to 100% mobile phase B over 7 min, and then 3 min at 100% mobile phase B. MS analyses were carried out using electrospray ionization in the positive ion mode using full scan analysis over m/z 200–1,100 at 70,000 resolution and 3 Hz data acquisition rate. Additional MS settings are: ion spray voltage, 3.0 kV; capillary temperature, 300°C; probe heater temperature, 300°C; sheath gas, 50; auxiliary gas, 15; and S-lens RF level 60.

We applied a previously published pipeline (https://github.com/broadinstitute/QC_metabolomics) to reduce noise in the profiling data and quantify differential metabolite abundances (Sevilla-Gonzalez et al., 2024; Sevilla-Gonzalez et al., 2022). The steps included normalization with internal standards and pooled samples, removal of metabolites with more than 25% missing values, imputation of missing values using half of the minimum detected value, Winsorization to ±5 standard deviations, and log2 transformation. Differential metabolite abundances between the two genotypes were then determined using Linear Models for Microarray and Omics Data (LIMMA) (Ritchie et al., 2015).

### EEG implantation and recording

To record electrical signals in the mouse brain, stereotaxic surgery (David Kopf Instruments) was performed on mice (n = 10 animals per genotype) to implant EEG electrodes (Pinnacle 8403, 0.10” mouse EEG screw with wire lead) intracranially above the frontal cortex (AP Bregma +1.5mm, ML Bregma +1.5 mm), parietal cortex (AP Bregma-1.5mm, ML Bregma +2.0mm), and a common ground and reference electrode above the cerebellum (AP Lambda-1.0, ML Lambda ±1.0). The electromyogram (EMG) electrodes were placed in the nuchal muscle of each mouse. Electrodes were soldered to an EEG/EMG headmount (Pinnacle 8201-SS) and encased in dental acrylic. Mice underwent surgery between the ages of 8-10 weeks and were allowed at least 2 weeks post-operative recovery according to the Broad Institute’s IACUC-approved protocol. Following surgery, mice were single-housed and allowed food and water *ad-libitum*. At 3-and 6-mo, mice were tethered to a Pinnacle EEG recording system (Pinnacle 8200-K1-SL) beginning with 48 hours habituation in single-housed recording chambers with food and water available *ad-libitum*. Passive EEG/EMG signals were then recorded for a full 24 hours during a 12/12-hour light/dark cycle. All signals were digitally sampled at a rate of 1,000 Hz, filtered 1-100 Hz bandpass for EEG, 10-1 kHz bandpass for EMG, and acquired using Sirenia Acquisition software (Pinnacle Technology).

### EEG data analysis

EEG recordings were divided into 10-second episodes, and each epoch was then staged as NREM, REM, or wake using Light Gradient Boosting Machine (LightGBM) algorithm (Wang et al., 2023b). Sleep spindles with different central frequencies (9, 11, 13, and 15 Hz) were detected using the Luna toolbox (Ghoshal et al., 2020) based on the determined threshold of WT animals. For power analysis, EEG power across different frequency bands, including delta, theta, alpha, sigma, beta, and gamma, was calculated. Outliers were excluded using the interquartile range (IQR) method, retaining only values within the range of Q1 − 2IQR to Q3 + 2IQR. The statistical significance between genotypes and frequency bands was calculated using linear mixed model analysis, with p < 0.05 as the significance criterion (for example, spindle density ∼ genotype* frequency + subjectID, where ID is a random variable). We used this model since it accounts for repeated measures across different frequencies and provides a robust evaluation of the effects of independent variables. The statistical effect of genotype was assessed using a likelihood ratio test by comparing two models: one that included the genotype-frequency interaction term (e.g., spindle density ∼ genotype * frequency + subjectID) and one without it (e.g., spindle density ∼ frequency + subjectID). For the analysis of spindle parameters and power at each frequency band, only the model with the interaction term was used (e.g., spindle density ∼ genotype * frequency + subjectID).

### OFT

The detailed methodology is outlined in our previous study (Herzog et al., 2023). Briefly, *Zmym2* mutant mice and their WT littermates (n ≥ 10 animals per group) were monitored for 60 minutes using the SuperFlex Open Field system (40 cm × 40 cm × 40 cm; Omnitech Electronics, Inc., Columbus, OH). The animals’ movement trajectories were recorded using Fusion system software (Omnitech Electronics, Inc.).

### Home cage monitoring

The methodology was described in detail in our previous study (Song et al., 2024). Male and female mice aged 1-mo were transferred to DVC (Tecniplast) and housed individually for the duration of the experiment. Briefly, the DVC racks utilize 12 electrodes beneath each cage that generate an electromagnetic field to detect disturbances corresponding to the animal movement. The DVC Analytics web platform was employed to calculate locomotor activity indices, which were calculated by the percentage of electrodes (out of 12) detecting movement over each 1-minute interval. An index value of 1 indicates movement detected by all 12 electrodes, while a value greater than 1 means more than 12 detected movements. The experiment included 10 female and 16 male mice (5 females and 8 males per genotype group).

### Bulk RNA-seq analysis

FASTQ files generated from each sequencing experiment were aligned to a reference derived from the genome FASTA file and transcriptome GTF extracted from the CellRanger mm10 reference through a Nextflow V1 pipeline (https://github.com/seanken/BulkIsoform) (Di Tommaso et al., 2017). Quantification was carried out with Salmon (Patro et al., 2017) (version 1.7.0, with arguments-l A--posBias--seqBias--gcBias--validateMapping), based on a reference generated using the Salmon index command with genomic decoys. Quality control metrics were obtained via STAR alignment (Dobin et al., 2013) (version 2.7.10a) and Picard tools CollectRnaSeqMetric command (version 2.26.7). Differential expression analysis was performed between heterozygous mutant versus WT mice for each brain region and age using R package DESeq2 (Love et al., 2014) (version 1.34). The Salmon output was loaded using R package tximport (Soneson et al., 2015) (version 1.22), and genes with at least 10 counts across all samples were included in the analysis. Log2 fold change shrinkage was applied with the “normal” shrinkage estimator in DESeq2. To analyze transposable element expression in RNA-seq data, we first realigned the FASTQ files with STAR (version 2.7.10a) with greater allowance for multimapping reads (--outFilterMultimapNmax 100--winAnchorMultimapNmax 200). We then used the TEtranscripts software package (Jin et al., 2015) (version 2.2.3) to test differential expression of transposable elements. We input the re-aligned BAM files, the gene GTF from our reference, and the curated mm10 transposable element GTF from the TEtranscripts website, and we included flags--sortByPos--format BAM--mode multi.

### snRNA-seq analysis

For snRNA-seq, FASTQ files were generated from raw BCL files and then aligned to a mm10 mouse reference genome using the Cell Ranger pipeline (Zheng et al., 2017) (v6.1.2). For Cell Ranger count,--chemistry=SC3Pv3 was used, and--expect-cells was determined by nucleus counting under a microscope. Within each experiment (i.e. brain region and age), replicates were downsampled using Cell Ranger aggr.

For 1-mo STR, ambient RNA removal was performed using CellBender (Fleming et al., 2023) after downsampling with DropletUtils (Lun et al., 2019) (v1.14.2).--expect-cells was determined as above and--total-droplets-included was set to 27000.

Seurat (Hao et al., 2024) (v4.0.3) was used to further process and analyze the Unique Molecular Identifier (UMI) counts. Nuclei expressing fewer than 500 genes were removed. The data from remaining nuclei were log-normalized and scaled by a factor of 10000, then scaled with ScaleData to prepare for dimensionality reduction. Linear dimension reduction was performed by RunPCA on variable genes. The nuclei were then clustered using FindNeighbors (with 20 dimensions) and FindClusters and visualized using Uniform Manifold Approximation and Projection (UMAP) using 20 dimensions. Doublet identification was performed using Scrublet (Wolock et al., 2019) (v0.2.3), and nuclei above a threshold determined by the score distribution were removed, as well as small clusters that were labeled in majority as doublets. Major cell types were labeled by examining expression of marker genes, and neuronal nuclei were re-clustered and subtypes were labeled using Azimuth (v0.4.6). Cell type proportions were summarized and differences were statistically evaluated using Speckle’s (v0.0.3) propeller function (Phipson et al., 2022).

Pseudobulking was used for differential expression analysis. For each gene, counts were summed across all cells of each cell type for each replicate. For each cell type, this pseudobulk counts matrix was then processed using edgeR (v3.36.0) (Robinson et al., 2010), first filtering out lowly expressed genes using the filterByExpr function. Surrogate Variable Analysis (SVA, v3.42.0) (Leek and Storey, 2007) was used to identify significant unknown latent sources of noise. These were added as covariates in the differential expression model matrix. Differential expression between the mutant and WT replicates was then performed using the likelihood ratio test.

### GSEA

To perform GSEA, we ranked the results using the negative logarithm of nominal p values multiplied by the sign of log2FC, derived from bulk RNA-seq, snRNA-seq, and proteomics differential expression results. These values were analyzed using the fGSEA v1.3.0 package (Korotkevich et al., 2021) with gene set collections, including M2 and M5 v2023.1 from the Molecular Signature Database (Castanza et al., 2023), SynGO gene collections (released on 2023/12/01) (Koopmans et al., 2019), gene lists of ENCODE histone modifications and ENCODE/ChEA transcription factor target gene sets from Enrichr (Xie et al., 2021), and gene sets curated from the literature (see Supplemental Table 8). For proteins with multiple isoforms, the isoform with the highest spectral count was used for GSEA. Mouse gene symbols from transcriptomic and proteomic data were mapped to their human homologs using annotations from Ensembl’s BioMart data service (Durinck et al., 2009), allowing GSEA to be conducted with human gene symbols. All significant GO terms (FDR < 0.05) across datasets, including bulk RNA-seq, snRNA-seq, and proteomics, are listed in the supplemental tables.

### Data and software availability

The original mass spectra and the protein sequence database used for searches are being deposited in the public proteomics repository, MassIVE (http://massive.ucsd.edu). Raw bulk and single-nucleus RNA sequencing FASTQ files will be deposited in NCBI GEO database. All datasets will be publicly available upon manuscript acceptance. The RNA-seq data for *Grin2a*, *Adnp*, and *Pogz* mutant mice used for comparison were previously published (Cho et al., 2023; Farsi et al., 2023; Suliman-Lavie et al., 2020).

