## Supplemental Figures for "Epigenetic changes, neuronal dysregulation and metabolomic abnormalities in *Zmym2* mutant mice, a genetic model of schizophrenia and neurodevelopmental disorders"

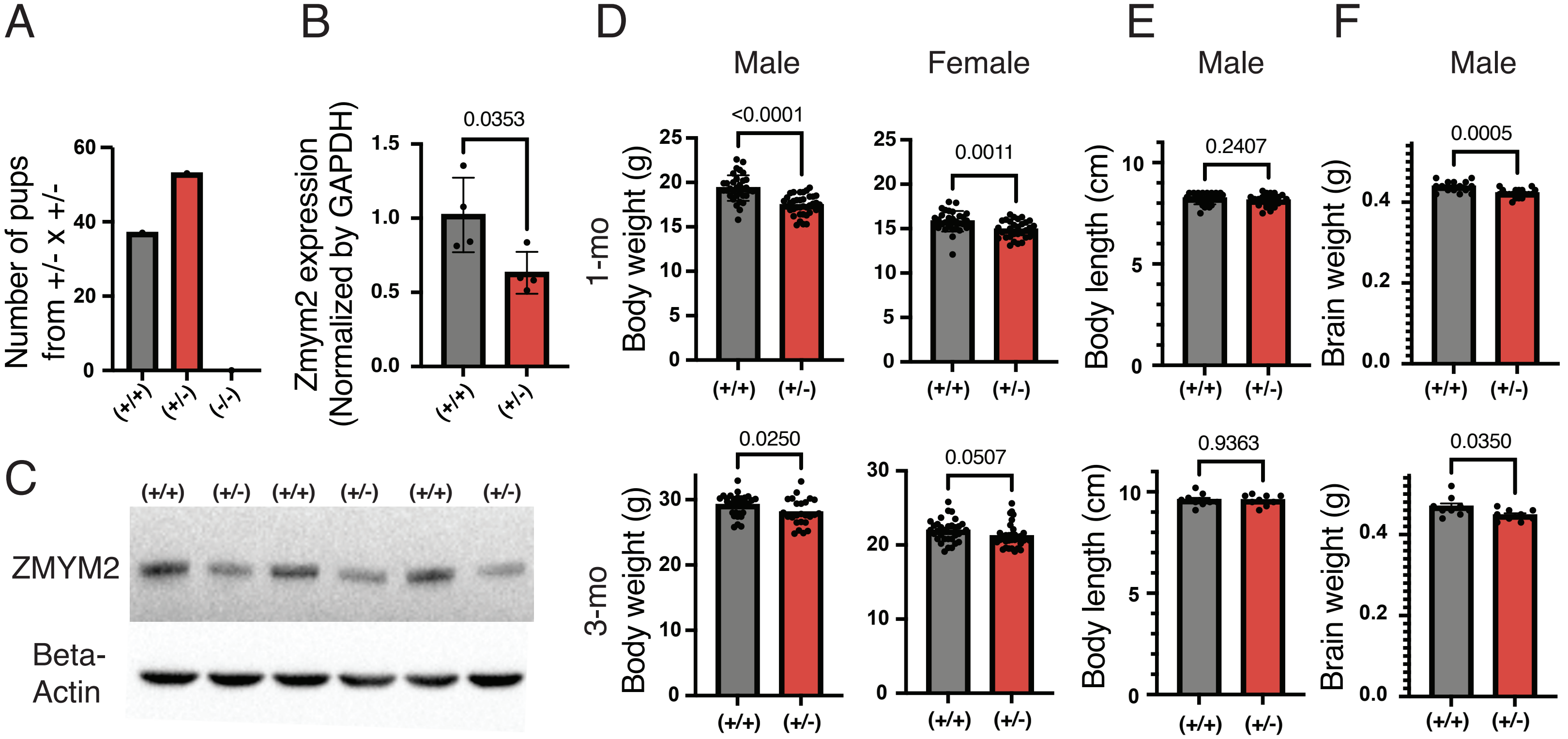


Suppl. Figure 1

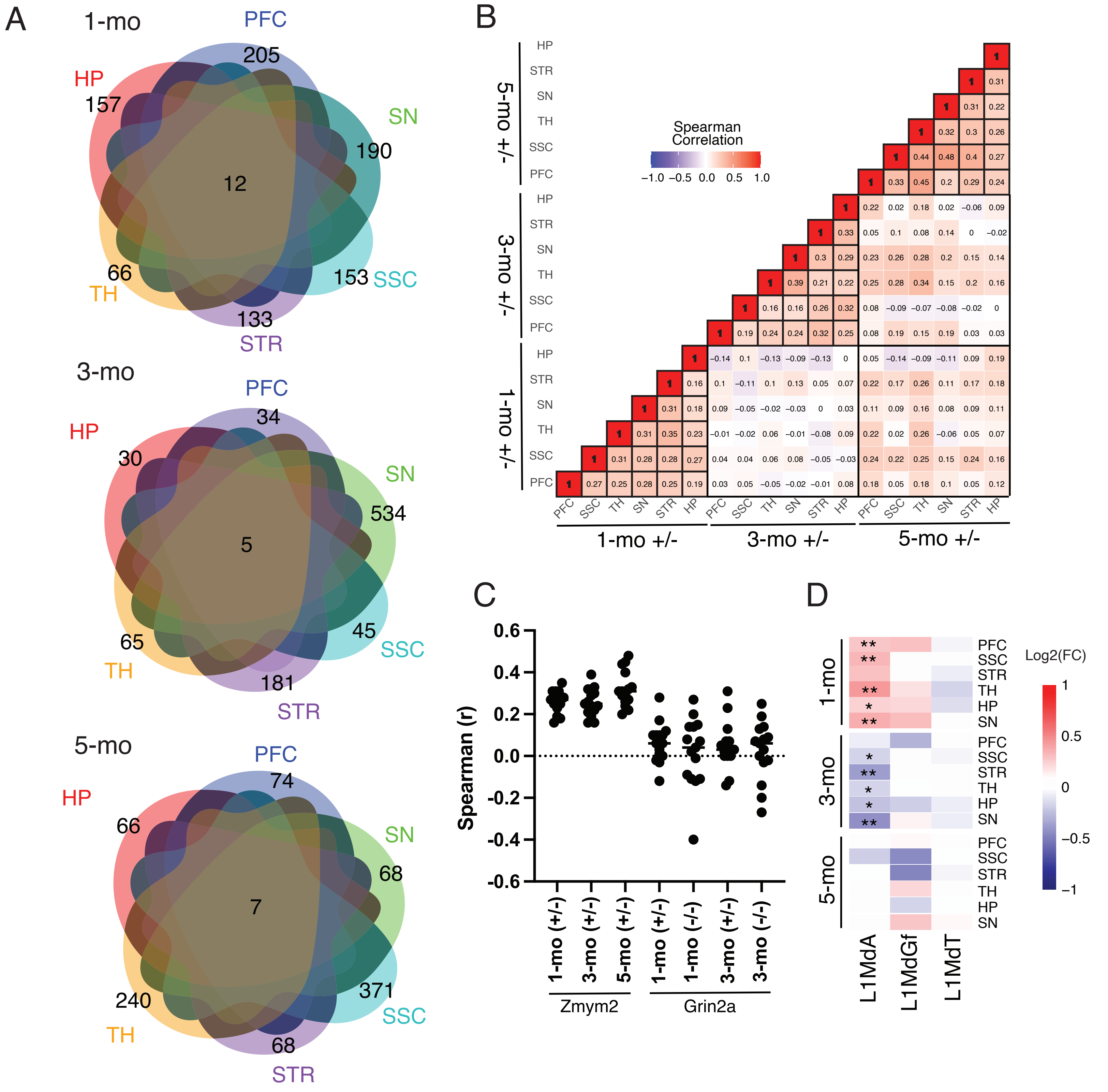


Suppl. Figure 2

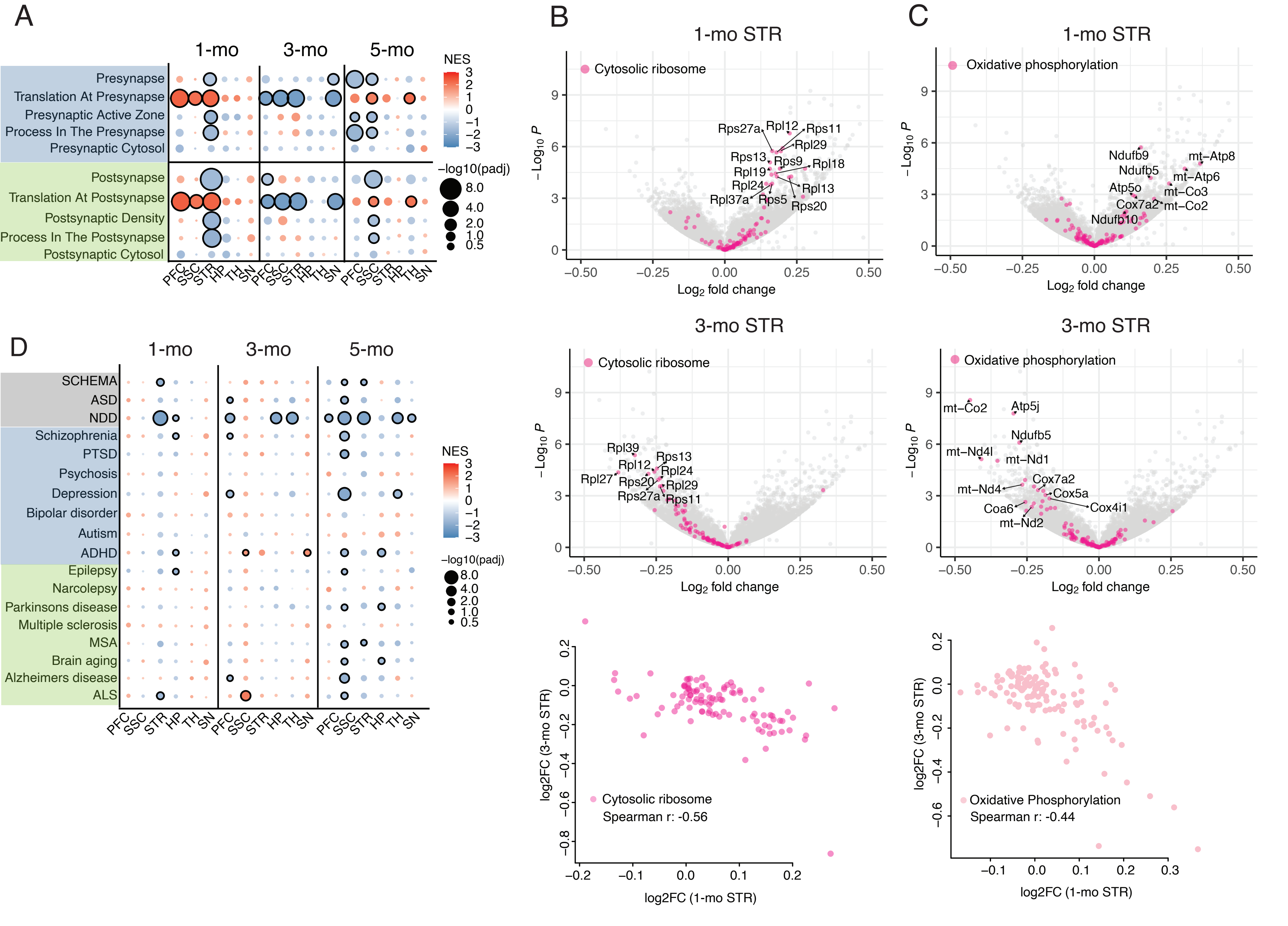


Suppl. Figure 3

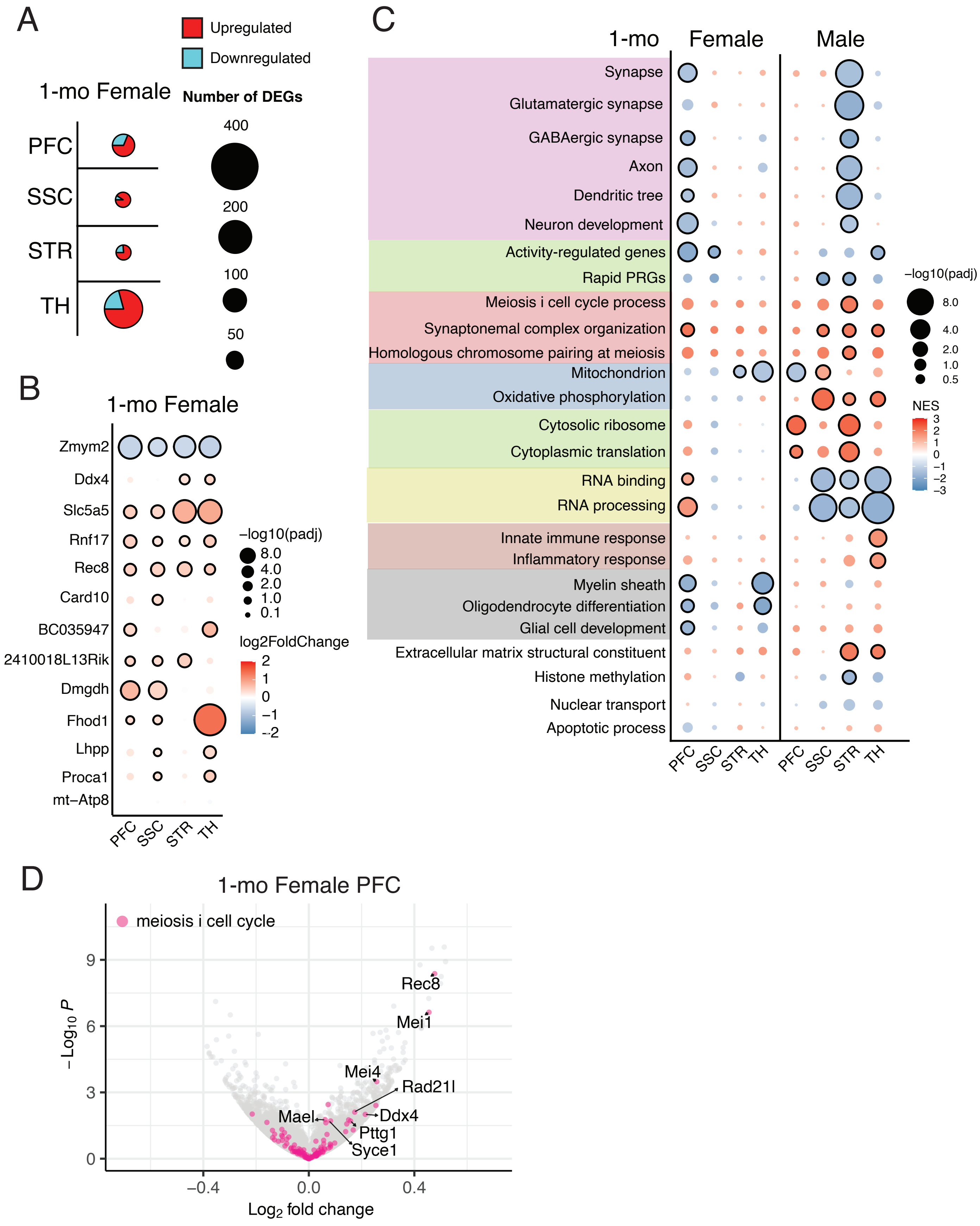


Suppl. Figure 4

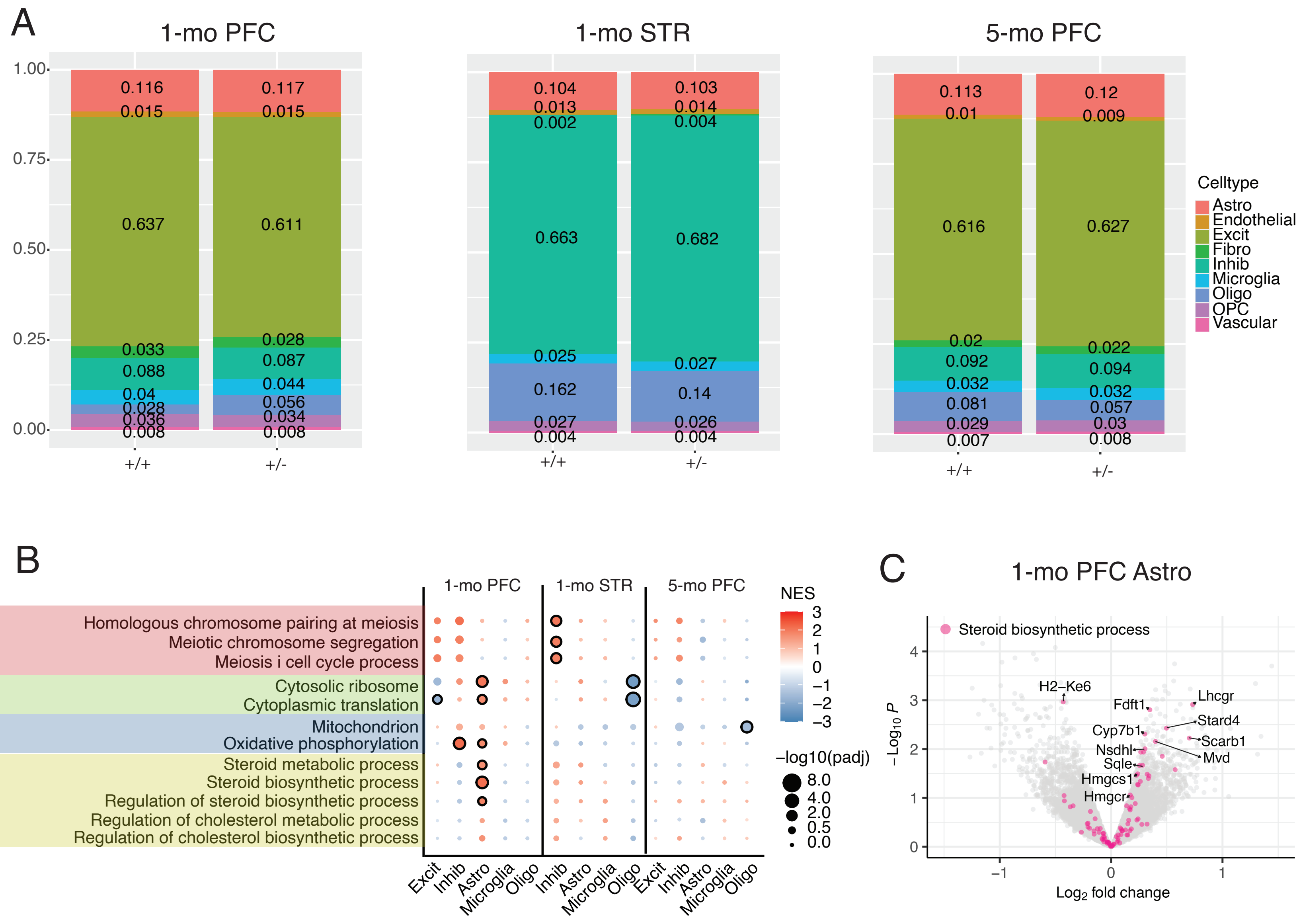


Suppl. Figure 5

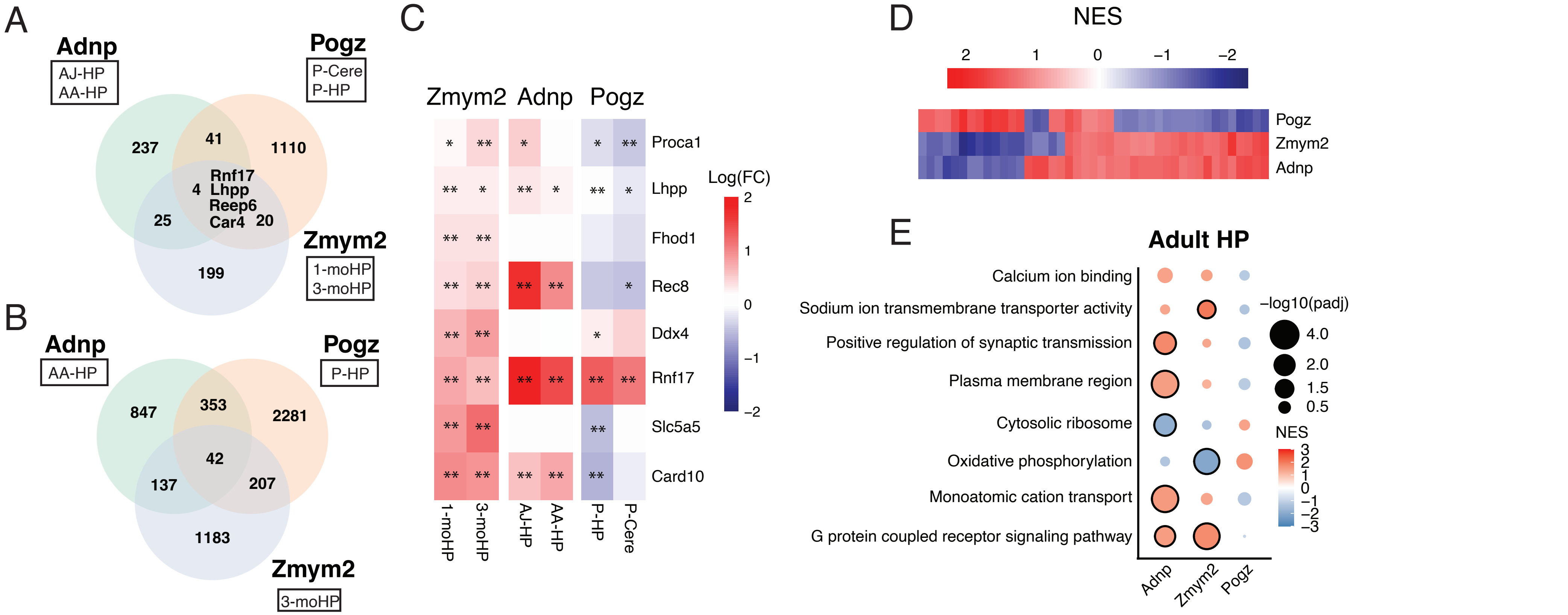


Suppl. Figure 6

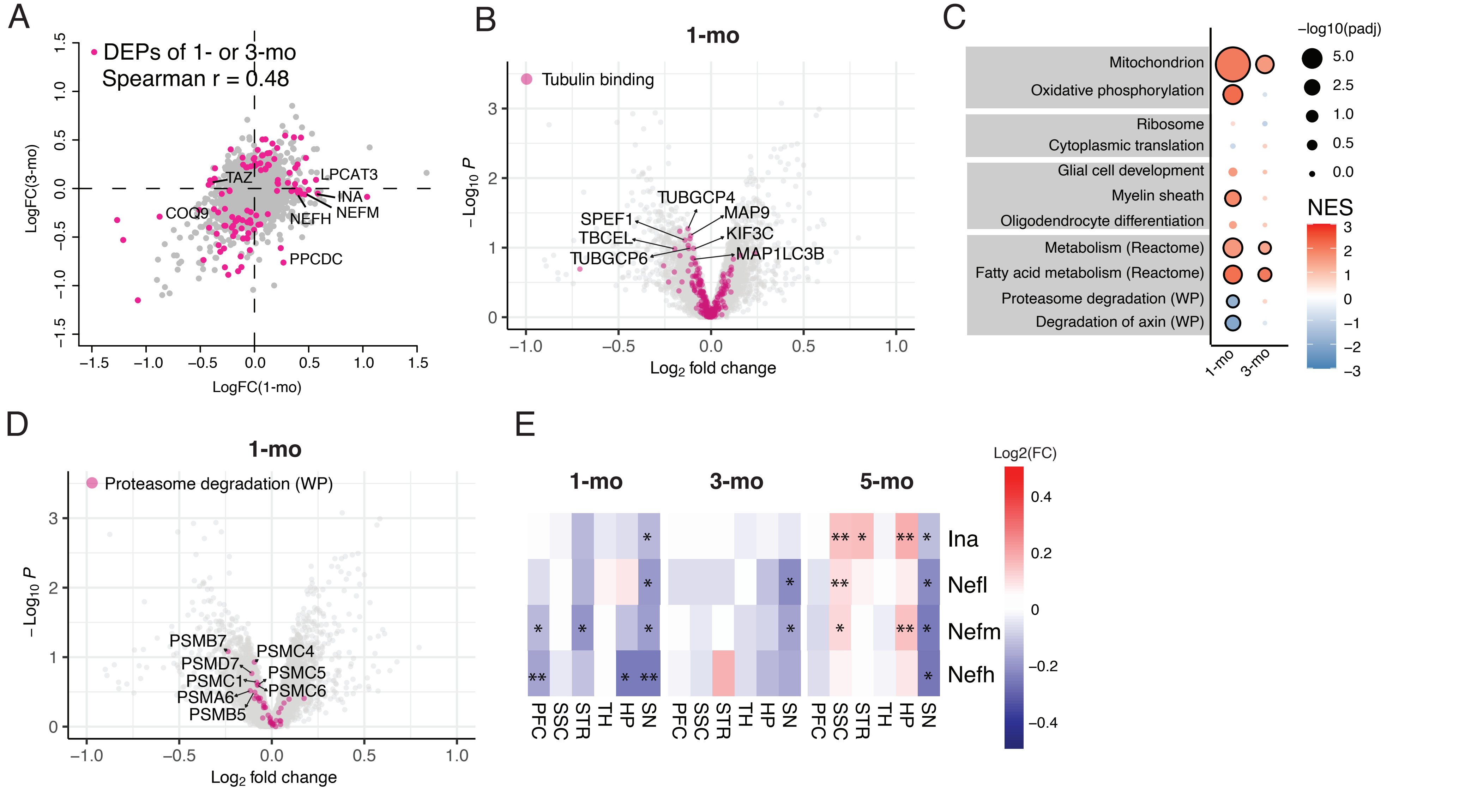


Suppl. Figure 7

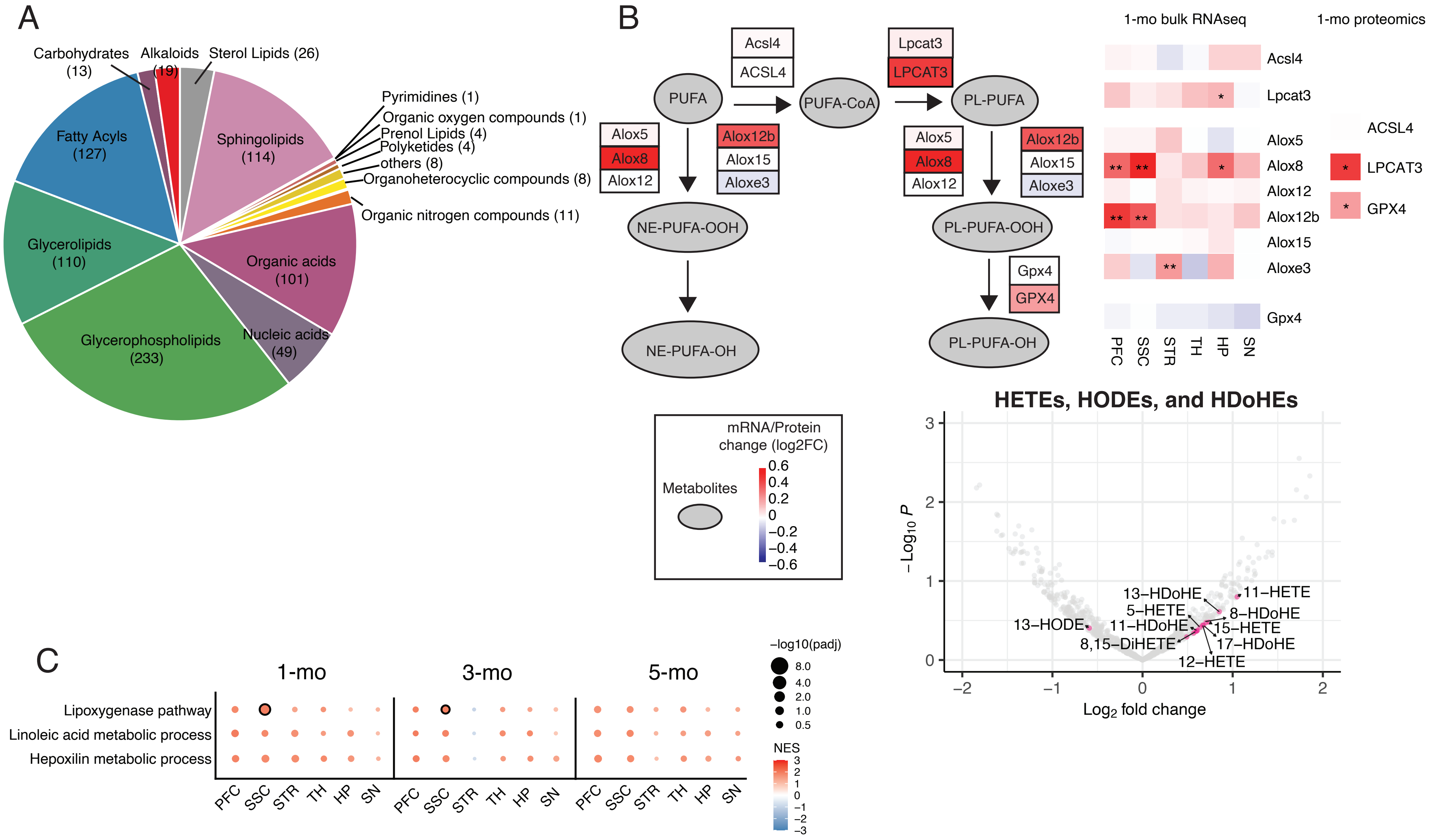


Suppl. Figure 8

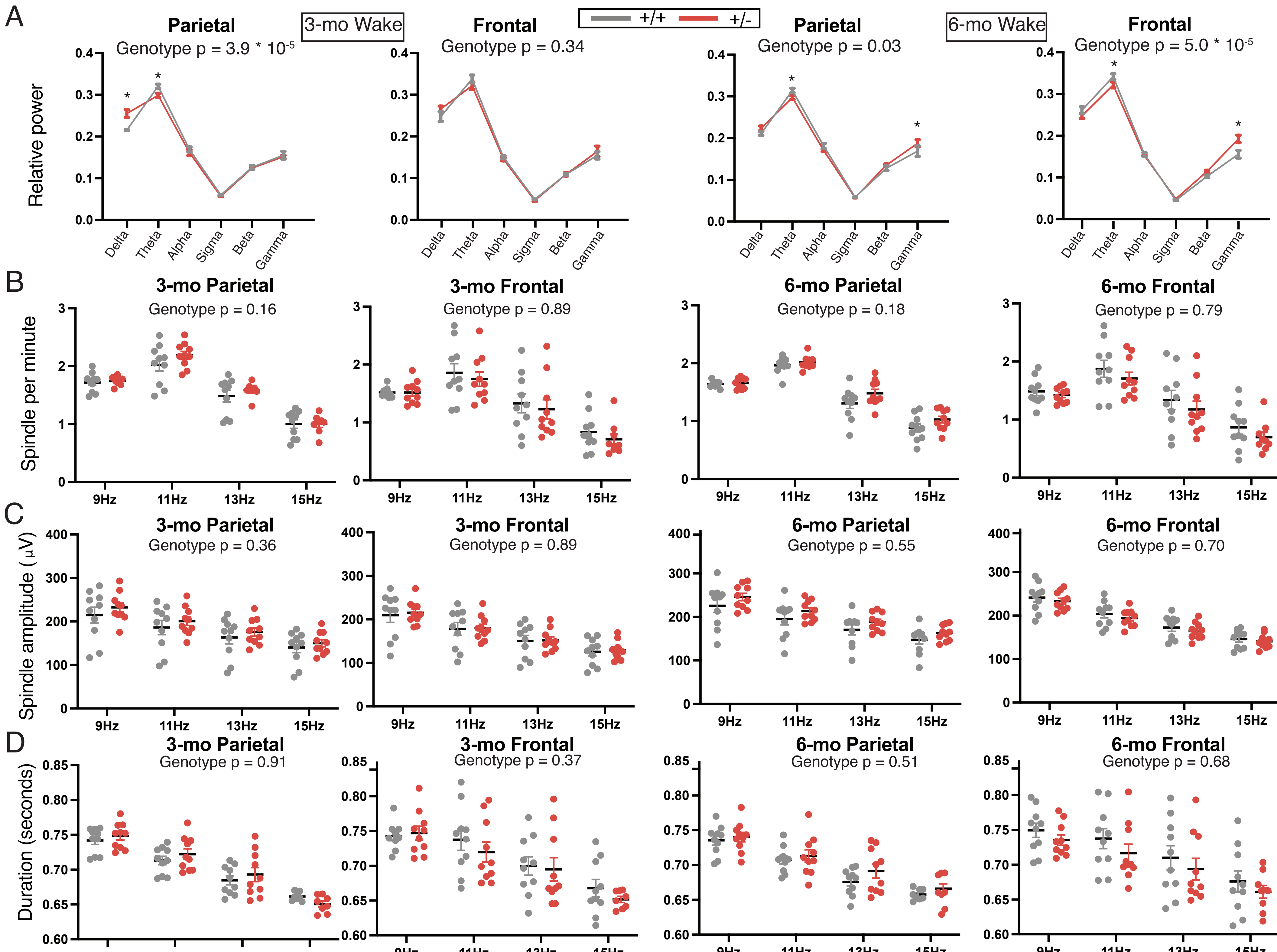

Suppl. Figure 9

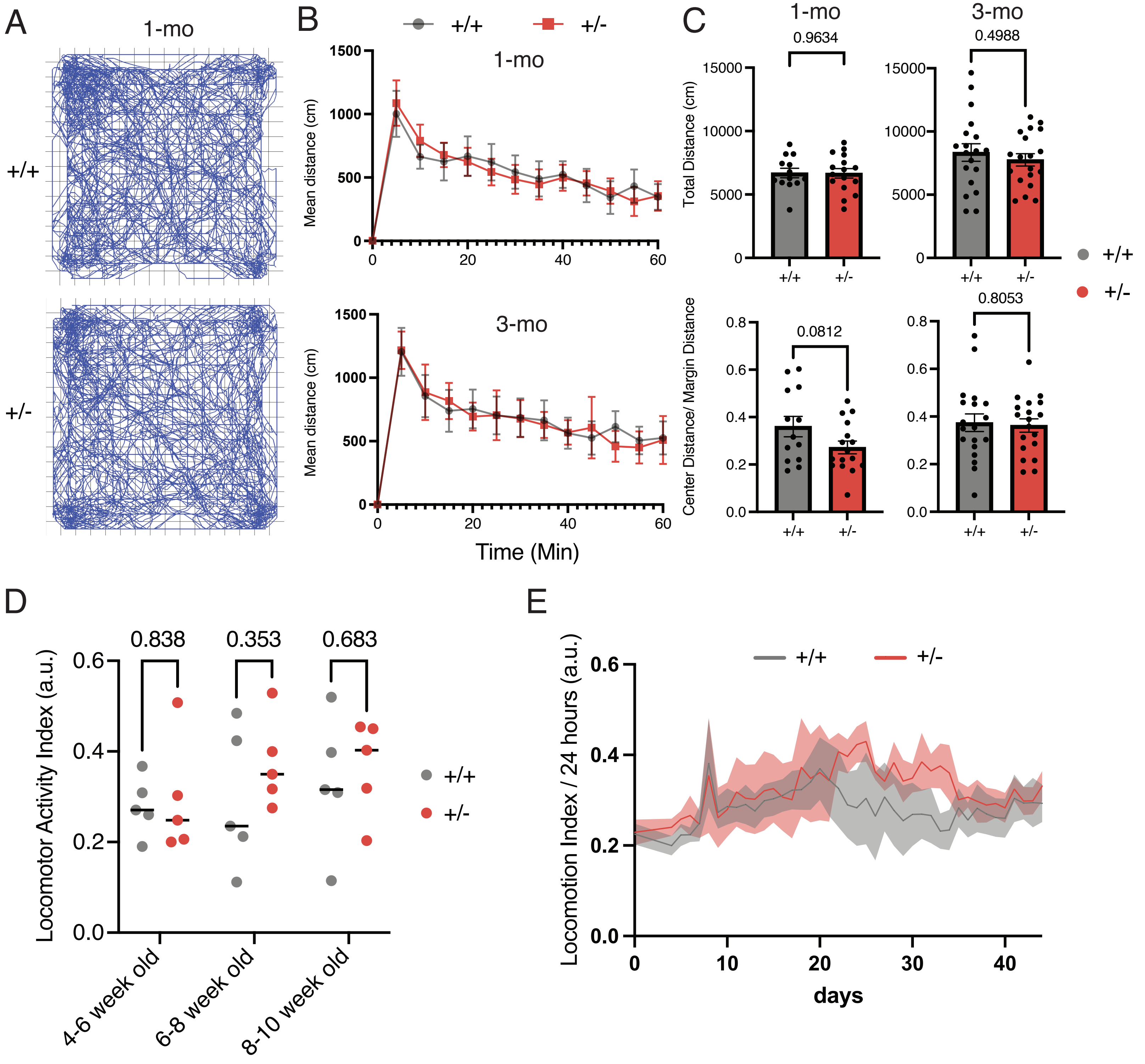


Suppl. Figure 10
